# The functional diversification of somatosensory neuron repertoires across Mammalia

**DOI:** 10.64898/2026.09.19.752904

**Authors:** S. Andrew Shuster, Jia Yin Xiao, Bruno Gegenhuber, Shamsuddin A. Bhuiyan, Min Dai, Yongting Chen, Michelle M. DeLisle, Preston Sheng, Elizabeth Howell, Michael Brecht, Elena O. Gracheva, Cynthia F. Moss, David L. Paul, Ishmail Abdus-Saboor, Gord Fishell, William Renthal, Michael E. Greenberg, David D. Ginty

## Abstract

The striking diversity of mammalian body forms and surfaces, lifestyles, and habitats prompts the question of how each species’ somatosensory neuron repertoire accommodates such diversity. The visual and olfactory systems diversify by gaining and losing sensor/receptor genes and corresponding sensory cell types. Using multiomic single-cell analysis of dorsal root ganglia neurons from thirteen different mammalian species and cross-species functional studies enabled by cell-type-specific enhancer viruses, we show that mammals instead assemble species-specific, functionally distinct somatosensory neuron repertoires from a conserved set of neuron types (orthotypes). Repertoire diversification takes multiple forms: orthotype abundance scales with variable bodily traits like hair follicle density, facilitating relevant behaviors; sensor/receptor expression is shuffled across species, such that orthotypes can detect different stimuli in different species; and orthotypes split into functionally distinct, species-specific subtypes (paratypes). Thus, unlike other sensory systems, the mammalian somatosensory system diversifies through multifarious changes to conserved orthotypes, enabling adaptation to diverse habitats, lifestyles, and body forms.

## Main Text

Natural selection shapes animal body structures and sensory systems alike to enhance survival and reproduction across diverse habitats. For instance, mammalian body surfaces vary widely, from the dense fur of the chinchilla, adapted to alpine Andean peaks, to the furless skin of the fossorial naked mole-rat, adapted to tunnels under the deserts and savannahs of East Africa. Many sensory systems, including the visual and olfactory systems, diversify by gaining and losing sensory receptor genes and corresponding sensory cell types which each express a single receptor (*1–8*). Somatosensory neurons transduce stimuli from across the body surfaces of all mammals, yet how mammalian somatosensory neuron repertoires have diversified to accommodate a wide range of environments, lifestyles, and body surfaces remains poorly understood.

Unlike other sensory systems, the mammalian somatosensory system is multimodal, coordinately detecting mechanical, thermal, and chemical stimuli, and many individual somatosensory neurons are themselves functionally polymodal, expressing multiple sensor/receptor genes simultaneously (*9–12*). This complexity has been dissected largely in mice: single-cell transcriptomic studies of mouse dorsal root ganglia (DRG) and trigeminal ganglion neurons (*11*, *13–20*) and functional studies using the vast mouse genetic toolkit have identified over a dozen molecularly, anatomically, and physiologically defined cell types (*11*, *12*, *17*, *21–32*). While this body of knowledge constitutes a foundation for defining mammalian somatosensory neuron types and their functions, the extent to which mouse somatosensory neuron types and their constituent features generalize across Mammalia remains unclear.

We asked how somatosensory neuron repertoires diversify across mammals by collecting and analyzing DRG neuron multiomic data from thirteen species spanning rodent, primate, and other placental clades, followed by cross-species functional studies enabled by new enhancer-based viral tools. We report that different mammals use a conserved set of neuron types (orthotypes) to assemble functionally distinct somatosensory neuron repertoires and show that these vary in three major ways: 1) orthotype abundance can scale with variable bodily traits like hair follicle density, facilitating relevant behaviors; 2) sensor/receptor expression is shuffled across species, such that orthotypes can detect different stimuli in different species; and 3) orthotypes split into functionally distinct, species-specific subtypes (paratypes). Thus, mammalian somatosensory neuron repertoires diversify not primarily by inventing new receptors and cell types, as other sensory systems do, but by recombining existing repertoires.

## Results

### Mammalian somatosensory neuron repetoires across rodent, primate, and other placental clades

To systematically describe how primary somatosensory neuron repertoires vary across Mammalia, we performed single-nucleus RNA- and ATAC-sequencing on DRGs harvested from all axial levels of nine species: four muroid rodents (mouse, rat, deer mouse, and hamster); two distantly related non-muroid rodents serving as an evolutionary outgroup (naked mole-rat and ground squirrel); and three small Laurasiatherian species (two bat species and Etruscan shrew) (Fig. 1A). We also re-analyzed publicly available DRG neuron single-cell datasets from guinea pig (*33*) and three old-world primates (two macaque species and human) (*33–35*). These 13 species belong to four clades that contain the most species-rich mammalian orders and ∼80% of all mammalian species (rodents ∼41%, bats ∼22%, Eulipotyphla ∼9%, primates ∼8%) (*36*). They collectively span lifestyles from human commensals (mice, rats) and hibernators (hamsters, ground squirrels) to fossorial (naked mole-rats) and flying (bats) mammals; body surfaces from hairy (most species) to nearly hairless (naked mole-rats and humans); and body masses from ∼2–3 g (Etruscan shrew) to 40+ kg (human).

**Fig. 1.**
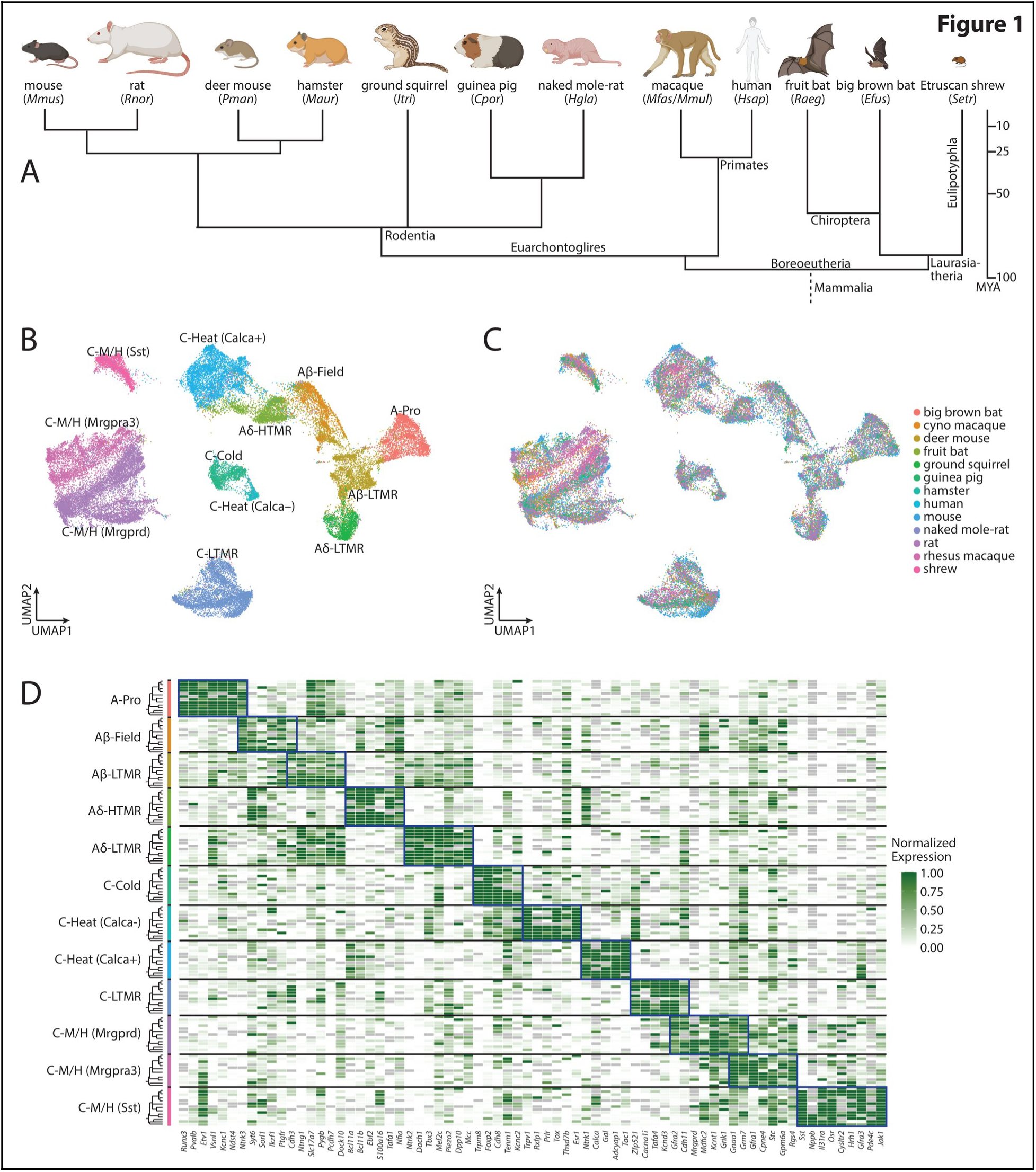
Mammalian dorsal root ganglia sensory neuron repertoires. (**A**) Phylogeny of mammalian species sampled. MYA, million years ago. (**B** and **C**) Uniform manifold approximation and projection (UMAP) embedding of single-nucleus RNA-sequencing integrated cross-species data. Colors indicate somatosensory neuron orthotype identities (B) or species-of-origin (C). (**D**) Heatmap showing average expression of marker genes (columns) within each orthotype across species (rows). Rows (left) are grouped by orthotype. Within each orthotype, species are ordered as in Figure 1A. Gray tiles indicate gene orthologs not present in any species annotation.

We used a standardized computational pipeline for quality control, normalization, dimensionality reduction, and clustering of each sample from each species separately before combining datasets across replicates and species for comparative analysis. Although our datasets contain non-neuronal DRG cells, our analyses focused on DRG neuron clusters.

### Conservation of DRG neuron types (orthotypes) across mammals

We first identified orthotypes of twelve major mouse somatosensory neuron types across species (Fig. 1 and fig. S1); ten of these twelve were conserved across all species, and only two species appeared to lack one or two orthotypes (see below). Large-diameter, heavily myelinated, fast-conducting A-fiber orthotypes included proprioceptors (A-Pro), which detect muscle, tendon, and joint movement and position (*37*); Aβ-Field, Aβ-, and Aδ-low-threshold mechanoreceptors (LTMRs) that detect light, innocuous mechanical forces (*11*, *24*, *28*, *31*); and A-fiber high-threshold mechanoreceptors (Aδ-HTMRs) that detect noxious mechanical and thermal stimuli and function as nociceptors (*11*, *12*, *31*, *32*). Small-diameter, slow-conducting C-fiber orthotypes included cold thermoreceptors (C-Cold) (*11*, *12*); nonpeptidergic putative warm (C-Heat (Calca–)) and peptidergic heat (C-Heat (Calca+)) (*11*, *12*) thermoreceptors; C-fiber LTMRs (C-LTMRs) (*24*); and three types of jointly mechano- and heat-sensitive (“nonpeptidergic”) polymodal C-fiber neurons (C-M/H neurons), which we distinguish here based on their mouse marker genes (*Mrgprd*, *Mrgpra3*, and *Sst*) (*11*, *12*, *25*, *26*, *38*).

Distinct analytical approaches relying either on conserved marker genes or on large numbers of highly variable genes all indicated widespread conservation of these orthotypes across most species. Cell types co-clustered across biological replicate samples from individuals of the same species (fig. S2, A to F), indicating high, consistent data quality. Combining samples from different species yielded graphs in which conserved cell types (orthotypes) co-clustered across species, suggesting conserved orthotypes across species and clades (Fig. 1, B and C). Many genes that mark specific DRG neuron types in mice had conserved orthotype-specific expression patterns across species (Fig. 1D) (*13*, *17*, *33–35*, *39*, *40*). In accordance with cell-type conservation, the expression patterns of several cell-type-restricted transcription factors (*17*) were largely conserved across orthotypes, as were those of growth factor receptors required for somatosensory neuron survival, growth, and cell-type identity (fig. S2, G and H) (*41*, *42*). Analysis of our snATAC-seq datasets likewise indicated conservation at the level of chromatin accessibility: orthotype-specific transcription factor binding-motif enrichment was largely consistent across species, suggesting that orthotypes use similar sets of transcription factors to establish and maintain core, conserved gene expression programs, and independent clustering of cells based on ATAC data recapitulated the neuronal clusters identified by transcriptomics (fig. S2, I to K). These transcriptomic and genomic data provide evidence of somatosensory neuron-type conservation across Mammalia.

Together, our single-nucleus RNA- and ATAC-sequencing datasets indicate that most of the DRG neuron types described and characterized in mice are broadly conserved across mammals. The near-complete conservation of somatosensory neuron orthotypes across a wide range of mammals suggests that each orthotype confers an adaptive advantage that has been maintained by selection. We provide an accompanying website (<u>link</u>) enabling conversion between different somatosensory neuron nomenclatures for easy access to these single-cell datasets as a community resource.

### Wide variation in DRG neuron orthotype proportions across mammals

Despite the remarkable conservation of somatosensory neuron orthotypes across most mammals, we observed marked variation in the proportions of several orthotypes across species (Fig. 2A and fig. S3). In a few extreme cases, an orthotype appeared to be absent in certain species. Most notably, C-LTMRs, which comprise ∼20% of all mouse DRG neurons, were absent in naked mole-rats and rare in humans (Fig. 2A and fig. S3). C-LTMRs are a large population of exquisitely mechanosensitive C-fiber neurons that respond to gentle stroking and cooling of the skin in mice (*24*) and have been implicated in pleasant/affective social touch (*43*, *44*), tissue repair (*45*), tactile allodynia (*46*, *47*), analgesia (*43*), pain recovery (*48*), anxiolysis (*46*), tickle (*49*), and wet-dog shake behavior (*50*). Moreover, studies of human C-LTMRs are inconsistent, with some assays describing sizeable C-LTMR populations (*51*) and others noting their rarity or absence (*35*, *51*, *52*). Given this apparent variability, we focused on characterizing cross-species differences in the proportions of this orthotype.

**Fig. 2.**
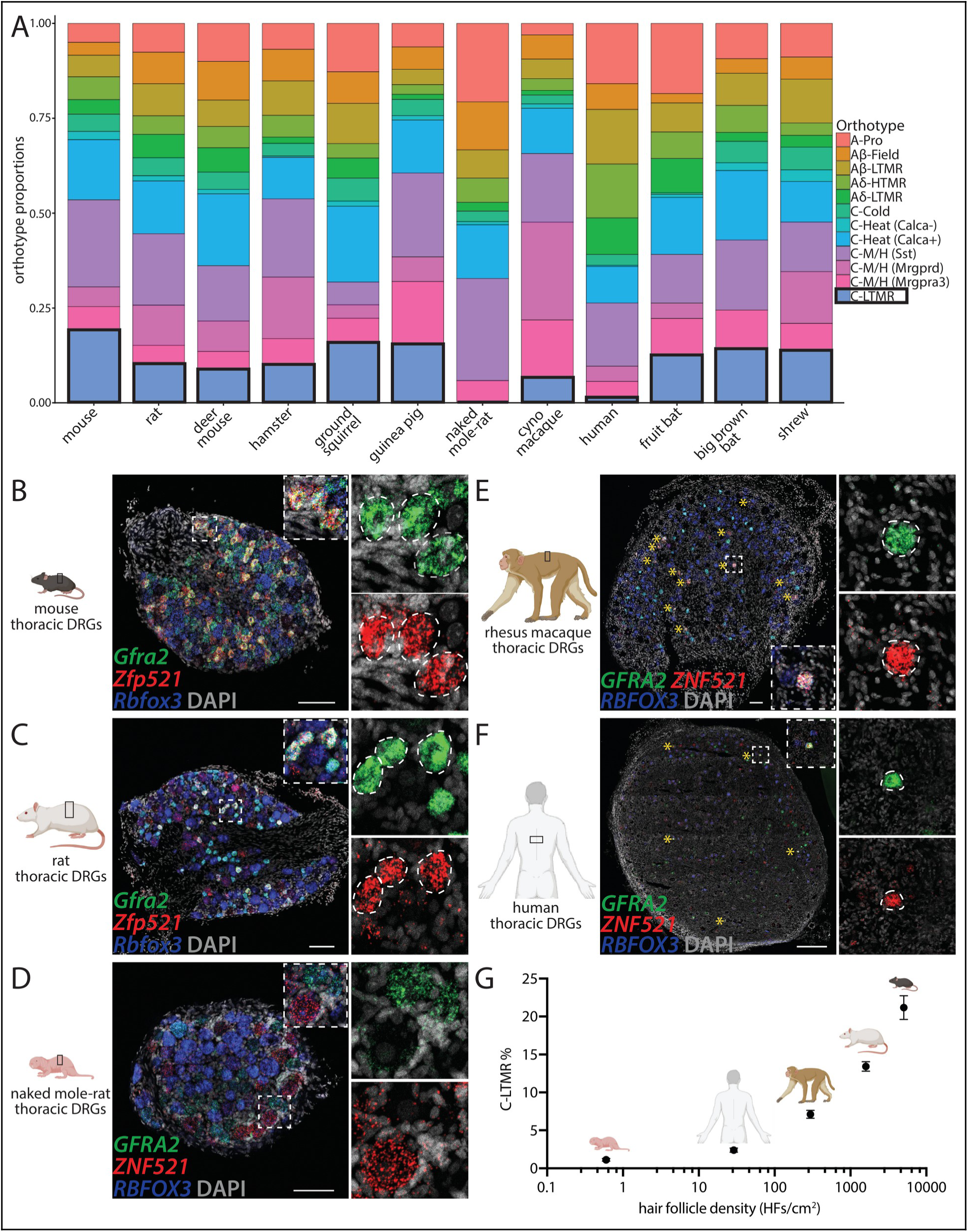
Variation across species in C-LTMR proportions in accordance with hair follicle density. (**A**) Averaged proportions of somatosensory neuron orthotypes across species. C-LTMR proportions are highlighted in black boxes (see also fig. S3). (**B** to **F**) Representative images of *in situ* hybridization against *Gfra2*/*GFRA2* (green) and *Zfp521*/*ZNF521* (red), co-markers of C-LTMRs, in mouse (B), rat (C), naked mole-rat (D), rhesus macaque (E), and human (F) mid-thoracic DRGs, along with pan-neuronal marker *Rbfox3*/*RBFOX3* (NeuN, blue). Putative C-LTMRs are outlined in dotted white lines in insets (right, all panels) and marked by yellow asterisks (E, F). Scale bars (in µm): 100 (B to E), 500 (F). (**G**) Relationship between hair follicle densities (see Methods) and C-LTMR proportions from mid-thoracic DRG sections; mean ± SEM (sections). Totals include 14–23 sections from 4–7 DRGs from 2–3 individual animals.

### Scaling of C-LTMR proportions and hair follicle density across mammals

C-LTMR proportions varied widely across species in our single-cell datasets: mice had the highest proportion (∼20% of all DRG neurons); most small mammals (rodents, bats, and shrews) had 9– 16%; macaques (∼7%) and humans (∼1%) had fewer; and naked mole-rat samples lacked C-LTMR clusters altogether (Fig. 2A and figs. S3 and S4, A to C). Notably, C-LTMRs are the only C-fiber neurons shown to innervate hair follicles, establishing lanceolate endings around the fine (zigzag and awl/auchene) hair follicles that comprise the downy undercoat and ∼99% of mouse trunk-skin hair follicles (*24*). C-LTMRs are also present at different proportions across axial levels in mice, with more at non-limb levels innervating hairy skin and fewer at limb-projecting levels (*17*, *24*). Given this intriguing relationship between hair follicle density and C-LTMR abundance in mice, we asked whether C-LTMR proportions also scale with hair follicle density across mammals, which exhibit several orders of magnitude in variation of this trait (*53*).

We performed *in situ* hybridization (ISH) against two conserved C-LTMR co-marker genes to quantify C-LTMR proportions in mid-thoracic DRGs, which innervate trunk hairy skin, across a range of mammals (mice, rats, naked mole-rats, macaques, and humans) with known hair follicle densities (Fig. 2, B to G and fig. S4, D and E) (*53*, *54*). Consistent with our single-cell data and previous studies (*17*, *24*), mouse mid-thoracic DRGs had abundant C-LTMRs (∼20%), proportionally more than those of rats (∼13%), which have a lower hair follicle density. By contrast, mid-thoracic DRGs of the naked mole-rat, a fossorial rodent lacking a downy undercoat (*54*), had exceedingly few (∼1%) putative C-LTMRs (Fig. 2D and fig. S4F). Assessment in DRGs of relatively hairy (rhesus macaque) and less hairy (human) primates (*53*, *55*) revealed far more C-LTMRs in macaques (∼7%) than in humans (∼2%) (Fig. 2, E to G). Thus, these data demonstrate that the relative abundance of an orthotype (C-LTMR) may scale with a salient, rapidly evolving mammalian body surface feature (hair follicle density).

### A C-LTMR-driven behavior conserved across hairy rodents

Motivated by the wide variation in C-LTMR proportions, we next asked what ethological functions C-LTMRs may play across mammals. C-LTMR functions have been assessed in mice largely by testing for deficits in candidate behaviors following C-LTMR genetic ablation or inhibition, and in humans by inferring cell-type identity from electrophysiological responses in peripheral microneurography studies while recording participants’ descriptions of evoked sensations (46, 56–58). We pursued a distinct gain-of-function approach, focusing on behavior resulting from stimulation of genetically identified C-LTMRs. C-LTMR stimulation in mice was recently reported to trigger wet-dog shake (WDS) behavior (*50*), a motor sequence in which animals repeatedly twist the head, neck, and upper trunk to forcefully expel water or other materials from the hair on their dorsal body surface (*59*). We therefore asked whether C-LTMRs innervate hair follicles and evoke WDS beyond mice, and whether naturalistic stimuli can evoke WDS in species lacking C-LTMRs.

Studying C-LTMR form and function beyond mice, wherein genetic tools are readily available, required new approaches. We therefore mined our single-nucleus multiome datasets to nominate ATAC peaks enriched in C-LTMRs as candidate enhancers and generated enhancer AAVs (eAAVs) (*60–65*) to test whether they could drive cell-type-specific payload expression across species (Fig. 3, A and B, and fig. S5A). Screening seven candidate elements revealed several that drove payload expression in mouse C-LTMRs with high cell-type-specificity (>75%; Fig. 3, C and D), including one located in an intron of a gene expressed specifically in rodent C-LTMRs (Fig. 3A and fig. S5, B and C). Importantly, this element also drove high, cell-type-specific reporter expression in rat C-LTMRs (Fig. 3, E and F), enabling cross-species comparisons of C-LTMR properties and functions.

**Fig. 3.**
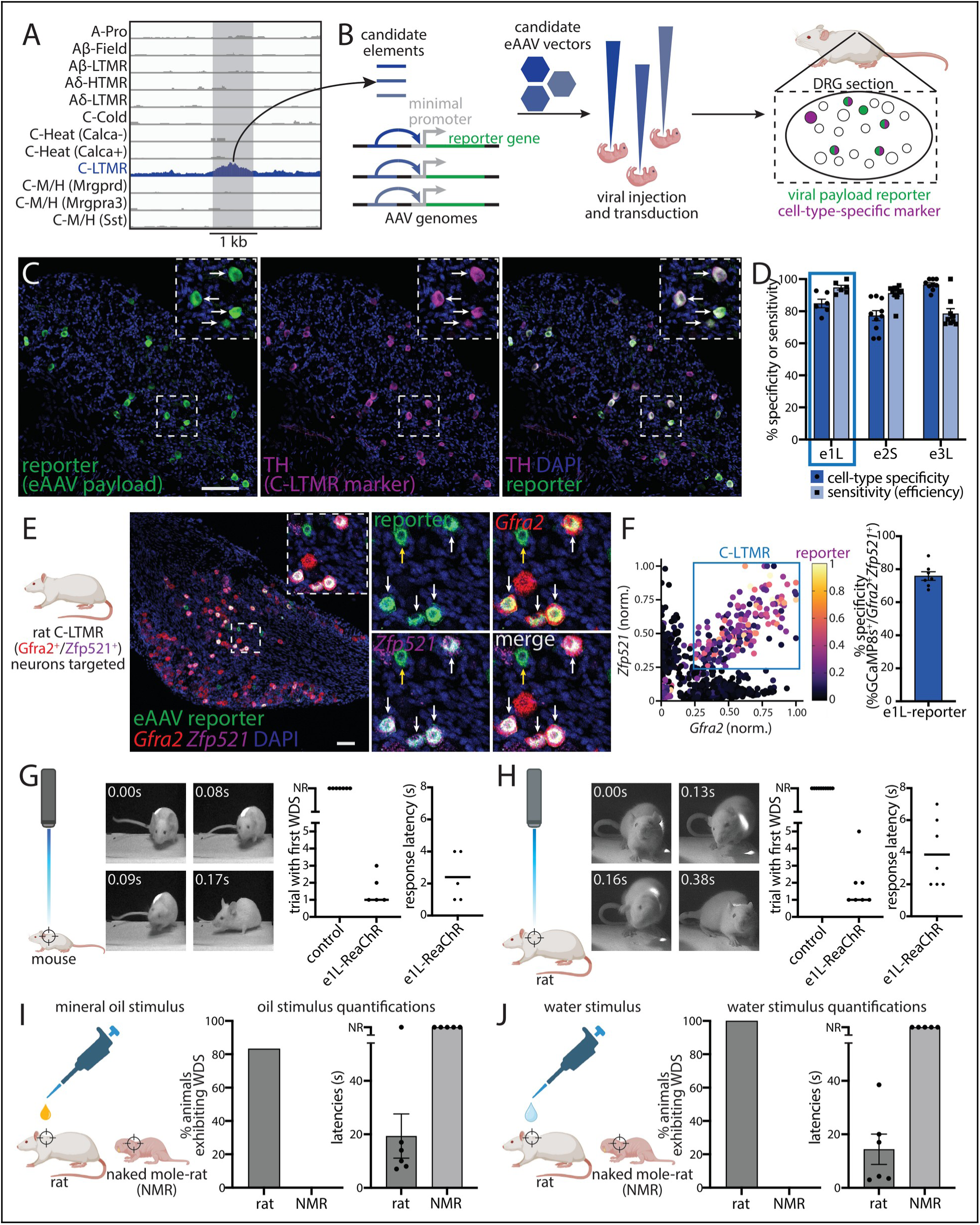
eAAV-enabled cross-species genetic access to C-LTMRs reveals control of conserved wet-dog shake behavior in hairy rodents. (**A**) Genome tracks showing a candidate cell-type-specific ATAC peak with enhanced accessibility in C-LTMR neurons (blue). (**B**) Schematic of *in vivo* eAAV screen. (**C**) Immunohistochemical validation of eAAV payload reporter expression in mouse C-LTMRs (TH^+^) (*24*). Insets, white arrows indicate TH^+^ neurons labeled by the eAAV reporter. Scale bar, 100 µm. (**D**) Quantification of specificity and efficiency of C-LTMR-targeting eAAVs in mice; mean ± SEM. Dotted line highlights eAAV tested and used in rats. (**E**) Joint *in situ* hybridization-immunohistochemistry validation of eAAV payload reporter expression in rat C-LTMRs (*Gfra2*^+^/*Zfp521*^+^). Insets, white arrows indicate *Gfra2*^+^/*Zfp521*^+^ neurons labeled by the eAAV reporter; yellow arrow indicates a *Gfra2*^–^/*Zfp521*^–^ off-target neuron labeled. Scale bar, 100 µm. (**F**) Quantification of specificity of C-LTMR-targeting eAAV in rats; mean ± SEM. (**G** and **H**) Left: schematics showing eAAV-mediated C-LTMR optogenetic stimulation of upper back. Middle: still frames showing mice (G) and rats (H) performing optogenetically induced wet dog shake (WDS) behavior and timecourse from WDS onset. Right: quantification of behavioral data in control (n = 7) and eAAV-C-LTMR-e1L-ReaChR (n = 5) mice and control (n = 10) and eAAV-C-LTMR-e1L-ReaChR (n = 7) rats. NR, none recorded. Bar, median trial with first WDS or mean response latency. (**I** and **J**) Left: schematics showing oil droplet (I) and water (J) application to upper back of rat and naked mole-rat (NMR) subjects. Right: quantification of behavioral data in rats (n = 6) and NMRs (n = 5). Oil droplets and water induced rat WDS behavior robustly, but never in NMRs. NR, none recorded. Latencies, mean ± SEM.

Using this eAAV, we first found that genetically labeled rat C-LTMRs, like those of mice, form hair follicle lanceolate endings and innervate spinal cord lamina II (fig. 6), indicating conserved peripheral morphology and central connectivity across these two species (*11*, *24*). We then asked if C-LTMR activation triggers WDS beyond mice. Optogenetic stimulation of the upper back in non-transgenic, virally injected mice and rats expressing an optogenetic activator targeted to C-LTMRs elicited WDS in experimental but not control animals (Fig. 3, G and H, Movies S1 and S2), revealing a conserved behavioral function across species (*50*). Further testing showed that rats, like mice, perform WDS following water or oil application to upper-back hairy skin (Fig. 3, I and J and fig. S7A). Because mice with C-LTMRs ablated show WDS deficits (*50*), we then asked whether species lacking C-LTMRs exhibit WDS in response to such stimuli. Strikingly, both water and oil applied to the upper back of naked mole-rats—which have few, if any, C-LTMRs and a fossorial lifestyle in which they rarely, if ever, get wet but constantly encounter debris on their dorsal skin (*72*, *73*)—failed to elicit WDS (Fig. 3, I and J and fig. S7B). Thus, furry C-LTMR-rich species (mice, rats, and macaques (*68*)) perform WDS, whereas species with sparse fur and few C-LTMRs (naked mole-rats and humans) appear not to (*59*). This is consistent with the proposal that WDS evolved to clear water trapped against the skin by fur (*59*), a need obviated in sparsely-haired mammals. Thus, the remarkable divergence of C-LTMR proportions across mammals corresponds to both hair follicle density and behavioral responses to hair-associated mechanical stimuli.

Together, our transcriptomic, anatomical, and behavioral analyses illustrate how diverse mammals employ conserved DRG neuron orthotypes in species-specific proportions adapted to their body surfaces, lifestyles, and habitats.

### Widespread sensor/receptor shuffling within orthotypes across species

While somatosensory neuron orthotypes can vary in abundance across species, they may also diverge in function if genes critical for sensory responses diverge in expression. We therefore asked whether DRG neuron orthotypes of different species across our datasets differ in the expression of ion channel and sensor/receptor genes. The expression of several ion channels that dictate cell-type intrinsic physiological properties (*69*) was largely conserved across species (fig. S8A). However, surprisingly, orthotype-specific expression of several critical sensor genes— including the mechanosensor *Piezo2* (*70*, *71*) and the thermo-/chemo-sensors *Trpa1*, *Trpm8*, and *Trpv1* (*72*)—diverged starkly across species (Fig. 4A), with numerous differences even between closely related species such as mice and rats. For example, mouse polymodal C-M/H (Mrgprd) neurons highly expressed *Trpa1* but not *Trpv1*, whereas the rat orthotype had the reciprocal pattern (Fig. 4A, rightmost green box). Many such differences were observed, especially within the three polymodal C-fiber (C-M/H) orthotypes. The Aδ-HTMR nociceptor orthotype also varied in *Trpm8* and *Trpa1* expression across species, suggesting potential species-specific mechanisms for detecting noxious cold and electrophilic substances. We call this phenomenon *sensor shuffling* and validated cases of it across several sensor genes (*Piezo2*, *Trpa1*, and *Trpv1*) in several mouse, rat, and human orthotypes by ISH (Fig. 4, A to J, and fig. S8, B to P).

**Figure 4.**
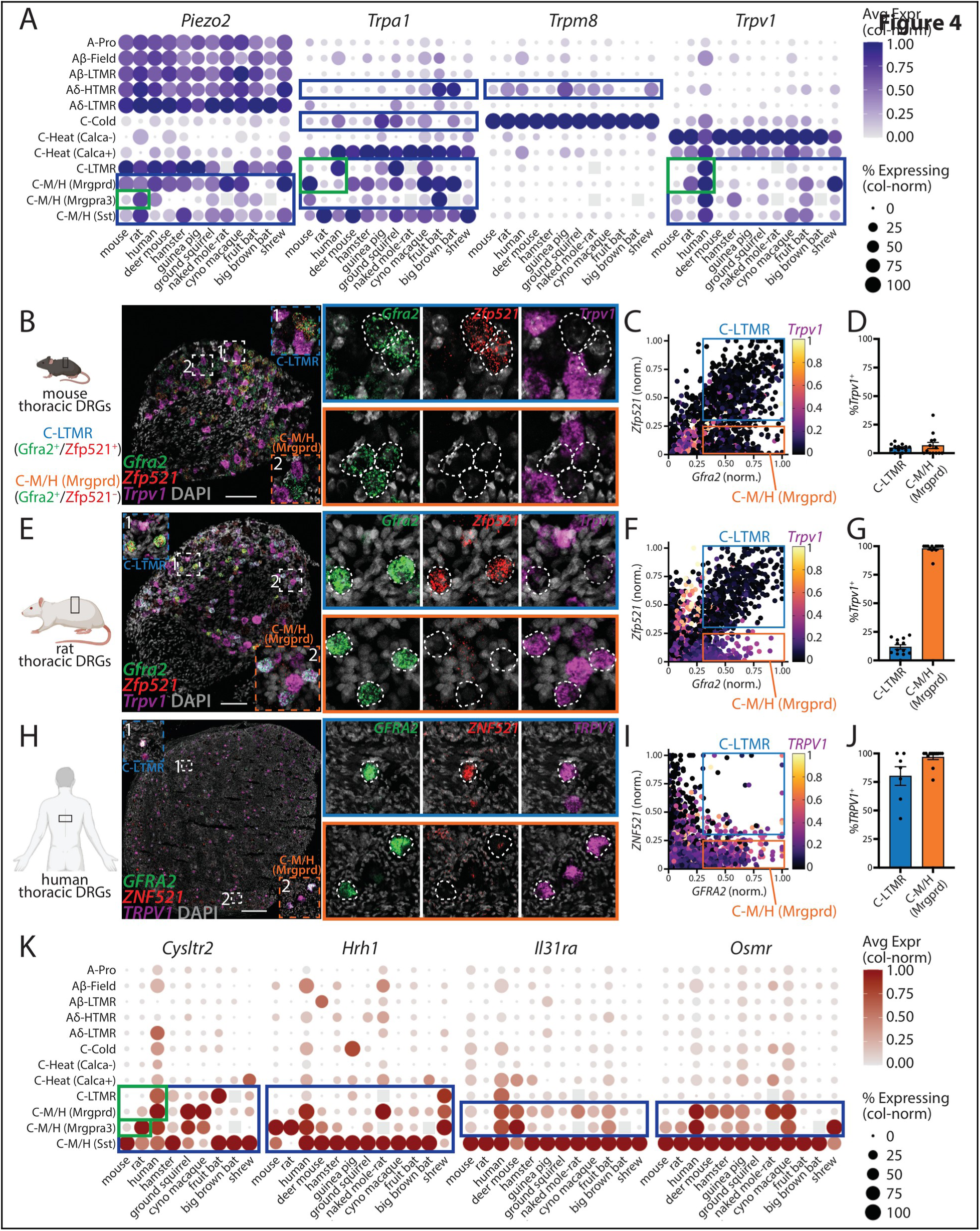
Conserved DRG neuron orthotypes exhibit dynamic sensor/receptor expression patterns across species. (**A**) Dot plot of sensor/receptor expression across conserved orthotypes (rows) and species (columns). Blue boxes indicate orthotype-sensor/receptor pairings exhibiting divergent sensor/receptor expression patterns (shuffling) across species. Green boxes indicate key sensor/receptor expression differences validated by *in situ* hybridization (see panels B to J and fig. S8). Gray boxes, missing cell types (see Fig. 2 and fig. S3). (**B** to **J**) *In situ* hybridization validation of *Trpv1* expression patterns in C-LTMR (*Gfra2*^+^/*Zfp521*^+^) and C-M/H (Mrgprd) (*Gfra2*^+^/*Zfp521*^–^) neurons across mouse (B), rat (E), and human (H) mid-thoracic DRGs. (C), (F), and (I): Quantification of neuronal *Trpv1* expression (2509, 3202, and 2439 neurons from 14, 13, and 12 sections from 6, 6, and 4 DRGs from 3, 3, and 2 mice, rats, and humans, respectively). (D), (G), and (J): Percentages of *Trpv1*^+^ neurons (14, 13, and 12 DRG sections from mice, rats, and humans, respectively); mean ± SEM. Scale bars, 100 µm (B, E), 500 µm (H). (**K**) Dot plot of immune receptor expression across conserved orthotypes (rows) and species (columns). Blue boxes indicate orthotype-receptor pairings exhibiting divergent receptor expression patterns (shuffling) across species. Green boxes indicate key receptor expression differences validated by *in situ* hybridization (fig. S10). Gray boxes, missing cell types (see Fig. 2 and fig. S3).

In some cases, orthotype-specific sensor expression patterns in mice (e.g., *Trpa1* and *Trpv1* in C-M/H (Mrgprd) neurons, *Trpm8* in Aδ-HTMRs, *Piezo2* in C-M/H (Mrgpra3)/C-M/H (Sst) neurons) more closely resembled those of deer mice or hamsters than of rats, the closest relative of mice in our datasets, ruling out phylogenetically constrained regulation of sensor expression. As these data indicate remarkable dynamicity in expression of these critical functional genes within and across orthotypes, even among muroid rodents, we asked how sensor shuffling between and across orthotypes emerges. Analysis of our snATAC-seq data suggested that, while cell-type-specific regulation of sensor expression by particular *cis*-regulatory elements can be conserved across species, orthotypes may also use distinct sets of *cis*-regulatory elements to regulate sensor expression (fig. S9), implicating divergent, species-specific distal *cis*-regulatory element (putative enhancer) utilization in the establishment of sensor shuffling.

This degree of sensor shuffling is unusual. Across many vertebrate sensory systems, notably vision and olfaction, sensory cells largely follow a one-receptor-one-cell-type rule (*2*), in which expression of a given sensor/receptor gene defines a cell type across species, and widespread inter-orthotype sensor/receptor gene shuffling has not been reported for other sensory modalities with conserved receptor genes (e.g., vision, taste). Somatosensory neurons appear to break this rule with their widespread sensor shuffling. Indeed, our findings suggest that sensor shuffling may endow orthotypes with flexibility of function across species.

This remarkable dynamism extends beyond canonical sensory receptors to receptors for internal, intercellular signals—notably those from immune cells, which DRG neurons detect alongside external stimuli. In fact, expression of immune/cytokine receptor genes was strikingly divergent across species (Fig. 4K). For example, *Cysltr2*, a receptor for immune cell-derived lipids (*73–75*), is a specific marker of mouse C-M/H (Sst) neurons (*11*) but was expressed in several other orthotypes across rats, ground squirrels, primates, bats, and shrews (Fig. 4K and fig. S10). Several other conserved immune/cytokine receptor genes (*Hrh1*, *Il31ra*, *Osmr*) likewise exhibited shuffled expression—including *Hrh1*, the histamine receptor implicated in histamine-mediated itch in humans (*76*, *77*), which our data show is expressed in different combinations of polymodal C-M/H orthotypes in mice, rats, and humans—suggesting that the corresponding immune signals are detected by different C-M/H orthotypes in different mammals. Notably, whereas immune receptor expression was largely restricted to the three polymodal C-M/H orthotypes in most rodents, some non-rodent C-LTMRs also expressed these receptors (*Cysltr2* in humans and fruit bats, *Hrh1* in shrew), suggesting that the ancestral genetic program specifying C-LTMRs retains the potential to confer immune signal detection, indicating a broader functional potential for C-LTMRs than previously appreciated.

The polymodal C-M/H orthotypes stood out in these analyses, exhibiting the most frequent shuffling of both sensors and receptors, while many other orthotypes showed largely invariant sensor/receptor expression. To ask whether this dynamicity reflects broader transcriptome-wide divergence rather than divergence of just a handful of genes, we built transcriptome distance trees across species and summed total branch length per orthotype (fig. S11) (*78*). A-fiber orthotypes showed, on average, lower transcriptomic divergence than C-fiber orthotypes, and the three polymodal C-M/H orthotypes were the most divergent overall, as they were with sensor/receptor expression. Together, these analyses suggest that the polymodal C-M/H orthotypes, which arise from a shared developmental lineage (*41*, *42*), are especially evolutionarily dynamic, both in critical sensor/receptor genes and at the whole-transcriptome level. Their extensive sensor/receptor shuffling also suggests that these orthotypes could play functionally distinct roles across species.

### Functional consequences of sensor shuffling across mouse and rat C-M/H (Mrgprd) neurons

Thus, to determine whether orthotypes with divergent sensor/receptor expression indeed have divergent sensory capabilities, we compared the physiological response properties of mouse and rat C-M/H (Mrgprd) neurons. Our transcriptomic and histological data indicated that mouse neurons highly express *Trpa1* but not *Trpv1*, while rat neurons highly express *Trpv1* but not *Trpa1*, predicting that mouse but not rat neurons would respond to the TRPA1 agonist allyl isothiocyanate (AITC) (*79*, *80*), whereas rat neurons would respond to the TRPV1 agonist capsaicin (*81*). Utilizing our multiome ATAC data, we generated eAAVs providing ≥85% (up to ∼97%) cell-type-specific access to C-M/H (Mrgprd) neurons in mice and rats (Fig. 5, A to E and S12, A to E), again demonstrating the cross-species utility of our eAAV platform. Labeled neurons of both species innervated spinal cord dorsal horn lamina II, indicating conserved central anatomy of this orthotype (fig. S13, A and B).

**Fig. 5.**
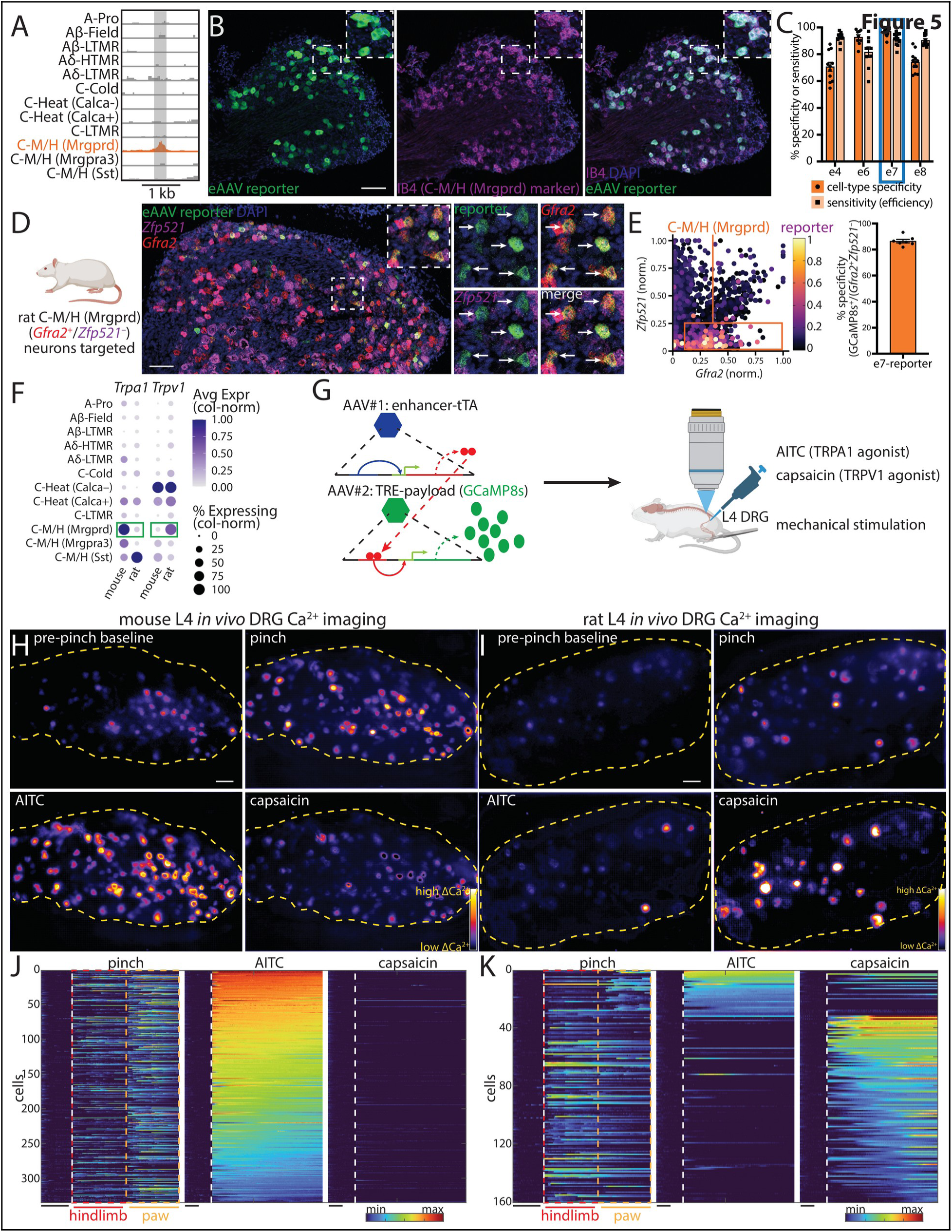
eAAV-enabled *in vivo* Ca^2+^ imaging in mouse and rat reveals C-M/H (Mrgprd) neuron orthotype functional divergence across species. (**A**) Genome tracks showing a candidate cell-type-specific ATAC peak with enhanced accessibility in C-M/H (Mrgprd) neurons (orange). (**B**) Immunohistochemical validation of eAAV payload reporter expression in IB4-binding mouse C-M/H (Mrgprd) neurons. Scale bar, 100 µm. See also figs. S12B and S12C. (**C**) Quantification of specificity and efficiency of C-M/H (Mrgprd) neuron-targeting eAAVs in mice; mean ± SEM. Blue box highlights eAAV tested and used in rats. (**D**) Joint *in situ* hybridization-immunohistochemical validation of eAAV payload reporter expression in *Gfra2*^+^/*Zfp521*^–^ rat C-M/H (Mrgprd) neurons. Scale bar, 100 µm. See also fig. S12A. (**E**) Quantification of specificity of C-M/H (Mrgprd) neuron-targeting eAAV in rats; mean ± SEM. (**F**) Single-cell RNA-sequencing data indicating opposite expression patterns of *Trpa1* and *Trpv1* in mouse and rat C-M/H (Mrgprd) neurons. (**G**) Schematic showing eAAV-mediated tTA-TRE-driven GCaMP8s expression in C-M/H (Mrgprd) neurons followed by the *in vivo* Ca^2+^ imaging preparation with mechanical and chemical stimulation. (**H** and **I**) Representative examples of *in vivo* Ca^2+^ imaging fields-of-view of pre-stimulus baselines (upper left) and responses (color bar) to pinch (upper right), AITC (TRPA1 agonist, lower left), and capsaicin (TRPV1 agonist, lower right) in mice (H) and rats (I). Yellow outlines, borders of DRG in field-of-view. Scale bars, 100 µm. (**J** and **K**) Summary heatmaps of Ca^2+^ responses to mechanical (pinch), AITC (TRPA1 agonist), and capsaicin (TRPV1 agonist) stimuli in mice (J) and rats (K). Data are sorted by responsiveness to AITC then capsaicin. N=350 cells, 9 DRGs, 7 mice (J) and 160 cells, 15 DRGs, 10 rats (K). Dotted white lines, stimulus onset. Colored bars, fluorescence (ΔF/F). Black bars, 10 seconds.

**Fig. 6.**
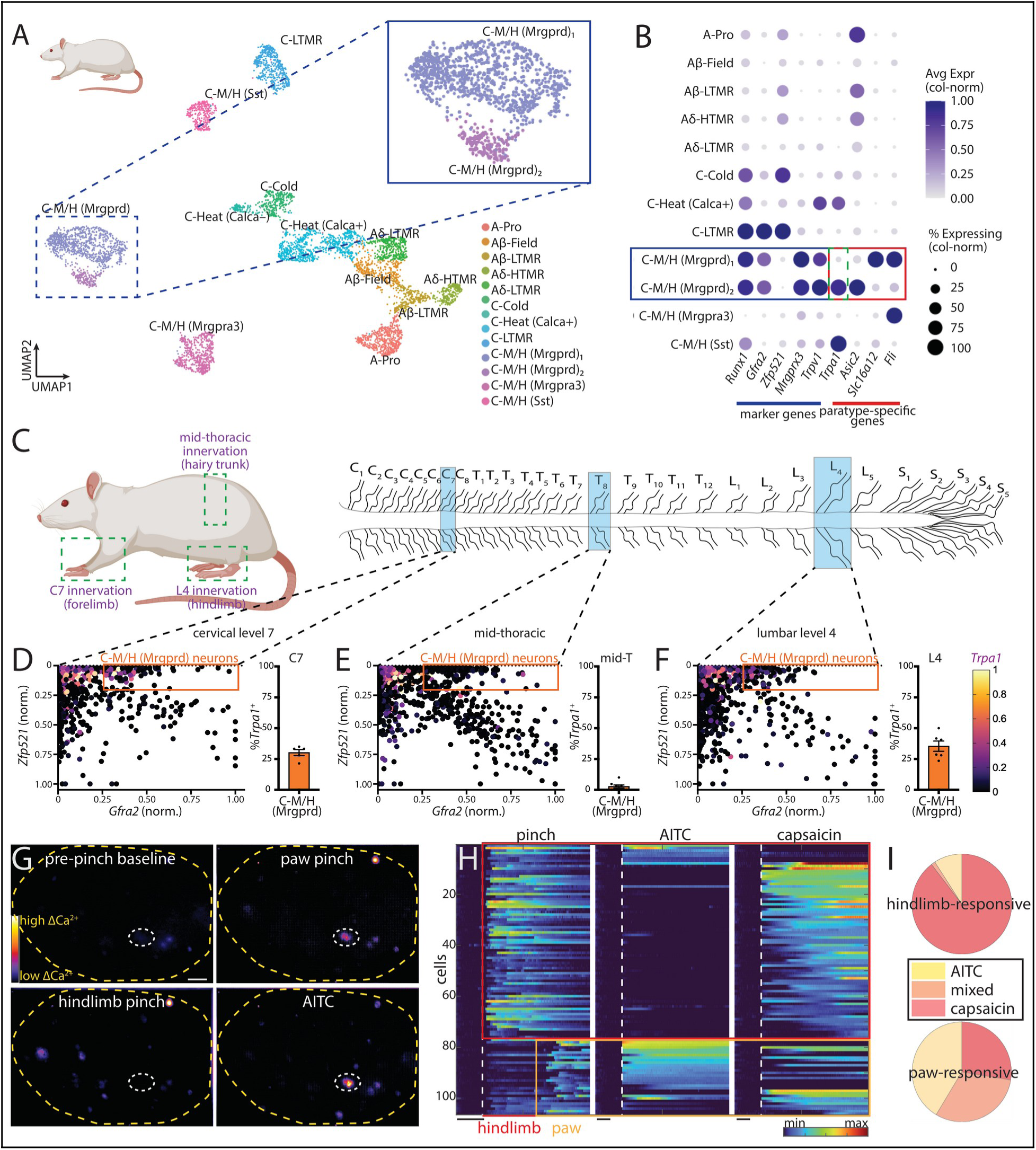
DRG neuron orthotypes diversify into anatomically and functionally distinct species-specific paratypes. (**A**) UMAP embedding of snRNA-seq data from one rat sample. Colors indicate somatosensory neuron orthotype identities. Inset shows splitting of C-M/H (Mrgprd) neuron orthotype into two putative paratypes. (**B**) Dot plot showing expression of common, orthotype-defining marker genes (blue line/box) as well as paratype-defining positive marker genes (red line/box), including *Trpa1* (dotted green box). (**C**) Schematic illustrating innervation of different peripheral cutaneous tissue types by DRGs from different axial levels, including cervical level 7 (C7), mid-thoracic levels (mid-T, e.g., T8), and lumbar level 4 (L4). (**D** to **F**) Quantification of neuronal expression of *Trpa1* by *in situ* hybridization (see fig. S14) in C-M/H (Mrgprd) and other neuron orthotypes in rat C7 (D), mid-T (E), and L4 (F) DRGs. Left, neuronal expression of *Trpa1* (color bar) across all quantified neurons. Right, percentages of *Trpa1*^+^ C-M/H (Mrgprd) neurons; mean ± SEM. (**G**) Representative images of *in vivo* Ca^2+^ imaging fields-of-view of pre-stimulus baseline (upper left) and responses (color bar) to paw pinch (upper right), hairy hindlimb pinch (lower left), and AITC (TRPA1 agonist, lower right). Note overlap between responsive cell signal (dotted white oval) across paw pinch and AITC stimulation. Yellow outline, border of DRG in field-of-view. Scale bar, 100 µm. (**H**) Summary heatmap of Ca^2+^ responses to hairy hindlimb (red box) and glabrous paw (yellow box) pinch-responsive neurons and their responses to AITC (TRPA1 agonist) and capsaicin (TRPV1 agonist). N = 106 cells, 15 DRGs, 10 rats. Traces are sorted by responsiveness to hindlimb pinch, then paw pinch, then AITC, then capsaicin. Dotted white lines, stimulus onset. Colored bar, fluorescence (ΔF/F). Black bars, 10 seconds. (**I**) Pie charts showing percentage of hairy hindlimb- and glabrous paw-responsive neurons that also respond to AITC (yellow), capsaicin (red), or both (orange).

We used a C-M/H (Mrgprd)-targeting eAAV to express the tetracycline transactivator to drive TRE-amplified GCaMP8s in mouse and rat C-M/H (Mrgprd) neurons and performed *in vivo* CaZ⁺ imaging of lumbar level 4 neurons to assess their physiological responses to mechanical and chemical stimulation (Fig. 5, F to K, and fig. S12, F to I). As predicted, C-M/H (Mrgprd) neurons of both species responded robustly to hindlimb mechanical stimulation. Remarkably, mouse neurons responded robustly to AITC but not capsaicin, whereas rat neurons responded robustly to capsaicin, with only a minor subpopulation responding to AITC (Fig. 5, H to K, and S13, C and D). These results establish that a conserved orthotype has been functionally repurposed across species through sensor/receptor shuffling and showcase that comparative transcriptomics can predict and cross-species cell-type-specific physiology enabled by eAAVs can reveal species-specific sensory functions at the level of individual orthotypes. More generally, these findings validate our single-cell transcriptomic results suggesting that sensor/receptor shuffling endows somatosensory neuron orthotypes with flexibility of function across species.

### Diversification of polymodal C-fiber orthotypes into species-specific paratypes

The minor AITC-responsive subpopulation of rat C-M/H (Mrgprd) neurons (Fig. 5, I and K, and fig. S13D) violated predictions based on the minimal *Trpa1* expression in the pseudobulk rat C-M/H (Mrgprd) dataset (Fig. 5F), despite cell-type-specific eAAV payload expression (fig. S12, H and I). Thus, we asked whether this AITC-responsive subpopulation corresponds to a species-specific C-M/H (Mrgprd) subtype—a paratype, in analogy to a gene paralog—that expresses *Trpa1*, which would indicate cross-species diversification of DRG neuron repertoires by species-specific orthotype splitting.

### Molecularly, anatomically, and physiologically distinct rat C-M/H (Mrgprd) neuron paratypes

Rat C-M/H (Mrgprd) neurons comprised two distinct clusters that we did not readily find in other species (Fig. 6A). While both expressed canonical C-M/H (Mrgprd) marker genes, the smaller cluster expressed *Trpa1* and other marker genes that the larger cluster did not (Fig. 6B), explaining the otherwise unexpected AITC responses, while the larger cluster expressed marker genes the smaller cluster did not. Given their shared expression of canonical orthotype markers, we interpret these not as de novo, species-specific neuron types but as candidate paratypes.

To distinguish true paratypes from mere cell states, we asked whether the clusters are anatomically distinguishable, performing ISH against orthotype and subcluster marker genes at different axial levels. One cluster was present at both limb- and non-limb-innervating levels, whereas the *Trpa1*-expressing cluster was present only at limb-innervating levels (Fig. 6, C to F, and fig. S14). Reexamination of our CaZ⁺ imaging data showed that most AITC-responsive neurons responded to hindpaw but not hairy hindlimb pinch, corroborating this anatomical distinction (Fig. 6, G to I). Together, these findings reveal diversification of rat C-M/H (Mrgprd) neurons into two molecularly, anatomically, and physiologically distinct paratypes and suggest that species-specific somatosensory neuron paratypes can specialize for innervation of distinct peripheral tissues. Paratype emergence may thus endow different skin types with distinct sensory capabilities.

### Paratype emergence across mammalian polymodal C-fiber neurons

Is such species-specific orthotype diversification into paratypes common? Our datasets revealed additional cases within the polymodal C-M/H orthotypes. Mouse C-M/H (Mrgpra3) neurons comprised two clusters not readily found in other species (figs. S15 and S16, A and B), corresponding to types annotated previously (*17*) and implicated in itch (*25*, *82*) and in pleasurable stroking touch and sexual behavior (*26*, *83*); while both expressed canonical orthotype markers, each also expressed paratype-restricted genes, including sensor/receptor genes. Datasets from distantly related non-rodent mammals also contained apparent paratypes: shrew C-M/H (Sst) neurons comprised two clusters differing in sensor and immune receptor expression (fig. S16, C and D). Together, these findings suggest that orthotypes conserved across many mammals—most frequently the polymodal C-M/H orthotypes—diverge into paratypes in a species-specific manner.

Thus, mammalian somatosensory neuron paratypes express not only shared, orthotype-defining marker genes but also collections of paratype-specific genes, including genomically conserved sensor/receptor genes that endow distinct sensory/functional capabilities. Somatosensory neuron paratypes can also be anatomically specialized, endowing different peripheral tissues with distinct sensory capabilities. Critically, these paratypes do not appear to have emerged in a sensory receptor gene paralog-linked manner, unlike in the olfactory and visual systems. This novel mode of receptor paralog-independent paratype emergence—a snapshot of orthotype diversification in action—suggests novel mechanisms of cell-type evolution for somatosensory neurons.

## Discussion

How has the mammalian somatosensory system changed with the radiation of mammals into diverse environments, lifestyles, and body forms and surfaces? Combining DRG neuron single-cell transcriptome and chromatin accessibility datasets from thirteen mammals with new enhancer-based viral tools for cross-species genetic access and functional analyses, we find that diverse mammals assemble distinct functional repertoires from a conserved set of orthologous somatosensory neuron types (orthotypes) that diversify in three ways: orthotype abundance, sensor/receptor expression patterns, and splitting into species-specific subtypes (paratypes). Together, these findings reveal that mammalian somatosensory neuron repertoires diversify across species not primarily by creating new cell types and sensor/receptor genes but by recombining existing cell types and sensor/receptor genes to meet species-specific needs.

### Conservation of mammalian somatosensory neuron orthotypes and tools for functional studies across species

Our transcriptomic and chromatin data indicate that the mammals profiled here assemble their somatosensory neuron repertoires from a conserved set of neuron orthotypes (Fig. 1 and fig. S2). This broad conservation distinguishes somatosensory neuron repertoires from those of other sensory systems, which diverge via genomic gain/loss of sensor/receptor genes and corresponding cell types (*2–7*). However, conservation of transcriptomically-defined cell types does not guarantee corresponding conservation of sensory function. Thus, leveraging our snATAC-seq data, we generated cell-type-specific eAAVs for cross-species genetic access enabling characterization of orthotype anatomy, physiology, and function. Because these tools readily permit payload customization and may be used in both transgenic and non-transgenic animals, they are useful both in mice and other species in which they have been validated. Furthermore, our snATAC-seq data and methods will enable development of a fuller viral toolkit for genetic access to mammalian somatosensory neuron orthotypes, including the orthotypes that initiate pain and itch signaling. Future comparative studies of cell-type diversity can follow the template demonstrated here, incorporating functional studies to test predictions generated by observational molecular/-omics datasets.

### Species-specific orthotype proportions

Although mammalian somatosensory neuron repertoires comprise largely conserved orthotypes, their proportions can vary greatly (Fig. 2 and fig. S3). Variation in sensory orthotype proportions has been noted in the retina (*8*, *84–86*), suggesting that cross-species variation in orthotype abundance may be a common evolutionary-developmental mechanism for tuning sensory systems to species-specific niches. The relationship we observe between C-LTMR proportions and hair follicle density suggests not only an intimate link between bodily surface features and somatosensory neuron development/specification, but also that hair-movement-associated mechanosensation may be a conserved C-LTMR function across mammals, complementing the diverse functions—notably affective touch (*46*, *87*)—previously proposed for this type. The sparsity of C-LTMRs at limb-projecting axial levels in mice (*17*, *24*), together with the relative rarity of this orthotype in human DRGs may also explain why single-cell RNA-sequencing studies of human DRGs, which often sample limb-innervating ganglia, variously report few or no C-LTMRs (*33*, *35*, *39*, *51*, *52*).

Differences in orthotype proportions appear in turn to constrain or enable particular behaviors, as we demonstrate for the C-LTMR-driven wet-dog shake (WDS): rodents with abundant C-LTMRs perform WDS, whereas naked mole-rats, which lack C-LTMRs, do not. Hair follicle density evolves rapidly in mammals (*61*, *93–97*) and may generally decrease with body size (*53*). Habitat, lifestyle, and thermoregulatory strategy could therefore jointly shape hair density, C-LTMR abundance, and the adaptive value of related behaviors like the WDS, which clears the dorsal body surface of liquids and irritants. An organism’s niche may thus shape the somatosensory neuron repertoire that organism assembles to meet the demands of its habitat, lifestyle, and body form and surface.

### Sensor/receptor shuffling in polymodal C-fiber (C-M/H) neurons

Because somatosensory neurons require sensor/receptor expression for sensory function, the widespread cross-species differences in their expression that we observe within orthotypes— sensor/receptor shuffling—imply divergence of orthotype function across species (Fig. 4), which our eAAV-enabled *in vivo* recordings confirmed for C-M/H (Mrgprd) neurons (Fig. 5). Sensor/receptor shuffling across species was most widespread within the three polymodal C-M/H orthotypes, which share central and peripheral anatomical (*11*), physiological (*11*, *12*), and behavioral (*13*, *25*, *38*, *82*, *93*) properties in mice. These orthotypes also express a range of immune receptors (*13*) and the evolutionarily labile Mrgpr chemoreceptor family (*94*) that detects both immune signals and environmental irritants (*95*, *96*). These commonalities suggest that these orthotypes may function redundantly and/or cooperatively, raising the question of why multiple such orthotypes are maintained. Because the functions of ancient multifunctional cell types tend to segregate among descendant sister types over evolution (*97*), it is tempting to speculate that the polymodal C-M/H orthotypes are more evolutionarily derived within mammals than other orthotypes, a possibility that could be tested by profiling distantly related species. Moreover, because immune-response genes are among the most rapidly evolving genes in mammals (*98*), the immune receptor shuffling largely restricted to these orthotypes suggests they may be rapidly modified across evolution and collectively tuned to detect diverse irritants and immune signals. Future testing of the functions of C-M/H orthotypes across species, perhaps using eAAVs, could also address such possibilities.

### Paratype emergence within polymodal C-fiber (C-M/H) orthotypes

We report that conserved orthotypes themselves diversify functionally by splitting into species-specific paratypes with distinct sensory capabilities, again most readily among the polymodal C-M/H orthotypes (Fig. 6 and figs. S14 to S16). While this cell-type diversification via paratype emergence is analogous to gene diversification by duplication-and-divergence, it does not appear to involve receptor paralog diversification, a critical feature of cell-type diversification in the olfactory (*1–4*) and visual (*5–7*) systems. Furthermore, partial functional redundancy among the polymodal C-M/H orthotypes may provide a permissive setting for paratype emergence, much as genetic redundancy permits paralog emergence following gene duplication. Moreover, the innervation of distinct peripheral tissues by functionally dissociable rat C-M/H (Mrgprd) paratypes suggests a link between peripheral targets and paratype development and thus a clue as to how somatosensory neuron paratypes could emerge in a receptor paralog-independent manner.

### Mechanisms of somatosensory neuron repertoire diversity across and within mammals

How the three forms of mammalian somatosensory neuron repertoire diversification identified here arise developmentally—whether specified intrinsically within post-mitotic lineages or instructed extrinsically by peripheral tissues—remains unaddressed; either mechanism could influence orthotype abundance through differential specification or survival. Axial-level (*17*, *24*, *35*) and dermatome-specific (*99*) (Fig. 6) patterning hints at an appreciable role for extrinsic instruction via retrograde peripheral signals. Intriguingly, the orthotypes with the greatest differences in proportions, sensor/receptor shuffling, and the most transcriptomic divergence across species (C-LTMRs and the three polymodal C-fiber orthotypes) all derive from a shared Runx1-dependent developmental lineage (*100–107*). As genes governing skin development and immune responses evolve rapidly (*98*), peripheral-tissue-directed programming of somatosensory neuron development could allow evolving skin and immune systems to flexibly tune corresponding somatosensory neuron repertoires during organismal development.

### Final Summary

In summary, mammalian somatosensory neuron repertoires assemble from a limited set of conserved orthotypes and diversify across species by three mechanisms—differences in orthotype abundance, sensor/receptor shuffling, and paratype emergence—each occurring both across species and across axial levels within species, and each most pronounced in orthotypes of a shared developmental lineage. By diversifying through such mechanisms, somatosensory neuron repertoires diverge across species in a way that other sensory systems do not, and in a manner that may be especially responsive to changes in peripheral tissues innervated by these neurons. Future work should address the developmental and evolutionary mechanisms generating these differences across Mammalia and within individual species.

## Acknowledgments

We thank members of the Ginty and Greenberg labs, especially X. Chen, T. Docter, E.E. Duffy, E. C. Griffith, D. Han, A. Lebedeva, K. Lezgiyeva, M. Lillis, S. Markman, R.I. Martinez-Garcia, K. Ng, J. Peng, A. Saha, C. Shi, J. Tycko, and X. Wu for discussions and comments on the manuscript. We also thank S. Hui and C.N. Rosique, S. Hrvatin, and H.E. Hoekstra for providing rat, hamster, and deer mouse samples, respectively.

## Funding

National Institutes of Health grant F32 (NS129589, S.A.S.)

National Institutes of Health grant K99 (NS146561, S.A.S.)

National Institutes of Health grant R35 (NS143029, M.E.G.)

National Institutes of Health grant R35 (NS132196/NS097344, D.D.G.)

National Institutes of Health grant R01 (NS119476, W.R.)

National Institutes of Health grant UO1 (MH142288, G.F.)

National Institutes of Health grant R37 (MH071679, G.F.)

National Institutes of Health grant DP1 (DP1AT013723, I.A.S.)

National Institutes of Health grant R01 (NS126271, E.O.G.)

National Science Foundation CRCNS grant (CBET-2011619, C.F.M.)

The Hock E. Tan and K. Lisa Yang Center for Autism Research (M.E.G., D.D.G.)

The Burroughs Wellcome Fund (W.R.)

The Rita Allen Foundation (W.R.)

The K. Lisa Yang Brain Body Center at Harvard Medical School (M.E.G., D.D.G.)

The Edward R. and Anne G. Lefler Center for Neurodegenerative Disorders (D.D.G.)

D.D.G. is an investigator of the Howard Hughes Medical Institute.

## Author contributions

S.A.S., M.E.G., and D.D.G. conceived the study. S.A.S. collected and analyzed single-nucleus data, with assistance from B.G. on ATAC data analysis and assistance from S.A.B. on cross-species integrative analysis. S.A.S., Y.C., and E.H. collected and analyzed histological data. M.D. ran the PIASO pipeline, with support from G.F. S.A.S. and Y.C. designed and performed eAAV screening, with assistance from D.L.P. on AAV production. S.A.S. designed, performed, and analyzed the Ca^2+^ imaging experiments, with assistance from J.Y.X. S.A.S. and M.M.D. designed, performed, and analyzed the mouse and rat behavior experiments. P.S. and S.A.S. designed and analyzed and P.S. conducted the naked mole-rat behavior experiments, with support from I.A.S. S.A.B. generated the website, with support from W.R. I.A.S., M.B., E.O.G., and C.F.M. provided samples. S.A.S., M.E.G., and D.D.G. wrote the paper with input from all authors.

## Data, code, and materials availability

Further information and requests for resources and reagents should be directed to David Ginty. All unique reagents generated in this study will be made available upon reasonable request. The original sequencing data will be deposited to the Gene Expression Omnibus (GEO: XXXXXXX) and made public upon publication of the manuscript.

## Supplementary Materials

### Materials and Methods

Figs. S1 to S16

References

Movies S1 to S2

## Materials and Methods

### Animals

All experiments performed in this study were approved by the Institutional Animal Care and Use Committee of Harvard Medical School. Experiments followed the ethical guidelines outlined in the NIH Guide for the care and use of laboratory animals (https://grants.nih.gov/grants/olaw/guide-for-the-care-and-use-of-laboratory-animals.pdf). C57BL/6 male and female mice (*Mus musculus*; C57BL/6J, The Jackson Laboratory, RRID:IMSR_JAX:000664) were used for sequencing and RNAscope in situ hybridization studies. CD1 male and female mice (Crl:CD1(ICR), Charles River, RRID:IMSR_CRL:022) were used for eAAV screening and eAAV-mediated physiological and behavioral experiments. Mice used for physiology and behavioral experiments were 4 – 8 weeks old. Sprague-Dawley male and female rats (*Rattus norvegicus*; Charles River, RRID:RGD_737891) were used for all rat experiments. Rats used for physiology and behavioral experiments were ∼3 and 4 – 8 weeks old, respectively. All mice and rats used for sequencing experiments were of weaning age (postnatal days 21 - 24). Roughly equal numbers of rodents of both sexes were used across all experiments. Mice and rats were housed in a temperature-controlled and humidity-controlled facility, maintained on a 12 h light/dark cycle, and given food and water ad libitum.

Tissue from rats (*Rattus norvegicus*; Charles River, RRID:RGD_737891), deer mice (*Peromyscus maniculatus bairdii*; Harvard University), hamsters (*Mesocricetus auratus*; Charles River/Massachusetts Institute of Technology, RRID:CVCL_TQ39), naked mole-rats (*Heterocephalus glaber*; Columbia University), ground squirrels (*Ictidomys tridecemlineatus*; Yale University), bats (*Rousettus aegyptiacus*, *Eptesicus fuscus*; Johns Hopkins University), and Etruscan shrews (*Suncus etruscus*; Humboldt University of Berlin) used for single-nucleus sequencing experiments were obtained in the laboratories of Drs. S. Hui (rat), H. Hoekstra (deer mouse), S. Hrvatin (hamster), I. Abdus-Saboor (naked mole-rat), E. O. Gracheva (ground squirrel), C.F. Moss (fruit bat, big brown bat), and M. Brecht (Etruscan shrew). Single-nucleus preparation was performed at Harvard Medical School for all samples except the naked mole-rat samples; single-nucleus preparation for these samples was performed at Columbia University.

### DRG Single-nucleus RNA- and ATAC-sequencing

We collected two to four biological replicates from distinct individuals for each species, including at least one male and one female whenever possible. Most samples were collected from healthy weanling or young adult animals. All rodent samples except the naked mole-rat samples were collected from healthy weanlings. DRGs from all axial levels were rapidly dissected out in DMEM-F12, HEPES (Thermo Fisher/Gibco, 11330-032; supplemented with 12.5 mM D-glucose and 1x pen/strep) on ice. DRGs were then snap frozen and stored at -80°C until further use or immediately processed for nuclear extraction and production of single-nucleus suspensions.

To produce DRG single-nucleus suspensions, DRGs were transferred to homogenization buffer (0.25 M sucrose, 25 mM KCl, 5 mM MgCl_2_, 20 mM Tricine-KOH, pH 7.8), dounced, and then layered onto OptiPrep gradients (OptiPrep™ Density Gradient Medium, Sigma-Aldrich, D1556-250ML). Various additives were included in homogenization and Optiprep/iodixanol solutions: 1 mM DTT (Life Technologies, 707265ML), 150 µM spermine-tetrachloride (spermine-tetrahydrochloride, Sigma-Aldrich, S1141-1G), 500 µM spermidine-trichloride (spermidine-trihydrochloride, Sigma-Aldrich, S2501-1G), protease inhibitor cocktail (Roche/Millipore Sigma, 11873580001; 100X stock made by dissolving two protease inhibitor tablets in 1 mL nuclease-free water), 10 mM sodium butyrate (Sigma, 19-137), and RNase inhibitor (Protector RNase inhibitor, Sigma, 3335402001; 1 U/µL for Multiome and 0.2 U/µL for 3’GEX). After ultracentrifugation (3,000 g for 15 minutes with no brake in a swinging bucket centrifuge at 4°C), the 30%–40% interface was collected, diluted and passed through a 40-µm filter. Nuclei were washed, pelleted, washed, and resuspended. Nuclei collected for Multiome (RNA and ATAC) samples were then processed for ATAC transposition, loaded onto 10x Genomics Chip J chips, and processed according to 10x Genomics Multiome protocol, while nuclei collected for 3’GEX (RNA only) samples were loaded onto 10x Genomics Chip G chips and processed according to 10x Genomics 3’GEX protocol.

10x Genomics protocols were followed to generate Multiome (RNA and ATAC) or 3’GEX (RNA only) libraries. Libraries were sequenced on NextSeq500, NovaSeq6000, or NovaSeq X Plus platforms, demultiplexed, and mapped to the appropriate reference genomes custom built using 10x Genomics pipelines. Reference genomes used for mapping included mm10 (mouse), mRatBN7 (rat), HU_Pman_2.1 (deer mouse), BCM_Maur_2 (hamster), HiC_Itri_2 (ground squirrel), Naked_mole-rat_maternal (naked mole-rat), mRouAeg1 (fruit bat), mEptFus1 (big brown bat), and mSunEtr1 (Etruscan shrew). Samples were sequenced to ≥19,979 mean reads per cell. After mapping to the appropriate reference genomes, we obtained new transcriptomes for a total of 59,200 (mouse), 108,194 (rat), 41,648 (deer mouse), 61,013 (hamster), 35,240 (ground squirrel), 66,507 (naked mole-rat), 86,131 (fruit bat), 72,450 (big brown bat), and 15,717 (Etruscan shrew) DRG nuclei, with a range of 15,717 – 108,194 cells across species and a mean of 60,678 cells per species. Mean reads per cell for each library: 51,327, 53,779, 38,565, 35,910 (mouse); 24,213, 27,899, 21,350 (rat); 31,141, 20,815 (deer mouse); 20,945, 24,850 (hamster); 25,296, 40,453 (ground squirrel); 26,132, 22,935, 19,979 (naked mole-rat); 23,763, 40,470, 23,262, 23,583, 49,672 (fruit bat); 26,026, 21,860 (big brown bat); 32,956, 90,527 (shrew). Median genes per cell for each library: 1,603, 1,228, 1,285, 801 (mouse); 1,274, 1,238, 1,358 (rat); 921, 745 (deer mouse); 1,695, 1,944 (hamster); 1,578, 707 (ground squirrel); 1,011, 833, 944 (naked mole-rat); 1,544, 477, 848, 1,161, 771 (fruit bat); 1,155, 951 (big brown bat); 822, 807 (shrew). Median genes per high-quality DRG neuron library were substantially higher than these per-cell averages (fig. S1). We integrated analysis of the above samples with analysis of samples previously collected from guinea pig (*Cavia porcellus*) and cynomolgus macaque (*Macaca fascicularis*) (*33*), rhesus macaque (*Macaca mulatta*) (*34*), and human (*Homo sapiens*) .

Data were processed using Seurat (*108*). Briefly, standard quality control metrics were applied. Data were then normalized, highly variable genes identified, and dimensional reduction and clustering performed. Clusters expressing non-neuronal gene markers were removed (*17*, *18*), leaving only neuronal populations. Clusters were annotated by expression of key conserved cell-type-specific genes (*13*, *17*, *33*, *34*, *39*, *40*) and validated using cross-species integration approaches.

### Single-nucleus Transcriptome Analysis

#### Standard quality control, analysis, and orthotype annotation

To extract high-quality neuronal data and determine the neuronal cell types constituent of each species’ DRG somatosensory neuron repertoires, we followed a standard quality control and cell-type annotation pipeline. For each sample, the single-nucleus RNA-seq count matrix was loaded from Cell Ranger output and initially filtered to retain high-quality nuclei. Gene expression was normalized using SCTransform, followed by Principal Component Analysis (PCA), Uniform Manifold Approximation and Projection (UMAP) embedding, and graph-based Leiden clustering (30 principal components, resolution 0.5); non-neuronal clusters were then removed and the normalization, dimensionality-reduction, UMAP embedding, and clustering steps were repeated on the retained nuclei, with canonical marker genes used to confirm neuronal identity. This process was repeated iteratively to obtain high-quality neuronal clusters (fig. S1).

#### Cross-species integration

To enable direct cross-species transcriptomic comparisons, scRNA-seq count matrices from all non-mouse species were first converted to mouse gene symbols. Ortholog mapping was predominantly executed using NCBI gene ortholog databases (109), supplemented with Ensembl BioMart exports for the naked mole-rat and guinea pig (110). Only 1:1 orthologs to mouse were retained for cross-species integration.

Downstream processing, clustering, and integration were conducted using Seurat. Quality control protocols filtered out low-quality cells, retaining only those with a minimum feature count (e.g., >1,500 detected genes) and a mitochondrial read percentage below 20%. Furthermore, human nuclei were specifically filtered to require MALAT1 expression greater than 2 (*111*). Following filtration, gene counts were log-normalized, and the top 2,000 highly variable features were identified independently for each species-specific dataset. To correct for batch and species-specific effects, datasets were integrated using Canonical Correlation Analysis via Seurat’s anchor-based integration framework. The integrated assay was scaled and subjected to PCA. Principal components (PCs) that cumulatively captured 90% of the variance, where no subsequent PC contributed more than 5%, were selected to construct a nearest-neighbor graph. Finally, cells were clustered using a graph-based approach (resolution = 3.0) and embedded into two-dimensional space using t-SNE and UMAP, with cell populations validated via the projection of established neuronal and non-neuronal marker genes. Clusters were reannotated based on marker gene expression profiles.

ShinyCell was used to build an interactive web resource to share the complete cross-species integrated object with the field (112).

#### Within-species biological replicate analysis

To test for consistency between biological replicates, we collected samples from distinct individual animals of the same species and merged the resulting datasets. Single-nucleus RNA-seq replicates of interest were loaded, labeled by their dataset of origin, and merged into a single object using raw RNA counts via the Seurat merge function. The merged, un-integrated data were then log-normalized, reduced to the top 2,000 variable features, scaled, and embedded by PCA and UMAP (30 principal components), and the resulting UMAP was visualized and colored by both replicate/dataset of origin and annotated with orthotype labels to assess data quality and consistency across replicates (fig. S2, A to F). Integration approaches were avoided to ensure high sensitivity to batch effects, technical noise, and/or biological variability.

#### Cross-species gene expression analyses (gene expression heatmap and dotplots)

For cross-species gene expression analyses, two major approaches to visualize data were taken. To display the relative expression levels of conserved orthotype marker genes, we generated a cross-species marker gene heatmap. For each sample, mean expression of the selected genes was computed per orthotype from the corresponding Seurat object, with gene names matched case-insensitively and genes absent from a given dataset recorded as missing values. Expression values were then min–max normalized to a 0 – 1 scale within each sample and gene and visualized as a heatmap of normalized expression across cell types and samples, with missing genes shown in gray. To display the relative expression levels of smaller numbers of sensor, immune receptor, transcription factor, growth factor receptor, and ion channel genes, we generated cross-species plots. For each sample, the selected genes (matched case-insensitively) were extracted from the corresponding Seurat object, and for each orthotype the mean expression and/or percentage of cells with detectable expression were computed. Within each sample and gene, both the mean expression and percent-expressing values were min–max normalized to a 0 – 1 scale across cell types and displayed faceted by gene, with point color encoding normalized mean expression and point size encoding the normalized percentage of expressing cells.

#### Orthotype whole-transcriptome orthology tree analyses

To ask if certain DRG neuron types are generally more variable across Mammalia, we sought to describe the relative rates at which the whole transcriptomes of different DRG neuron orthotypes have diverged rather than relying on the differential expression of a small number of sensor and receptor genes. To do so, we assembled transcriptome distance trees, a measure of transcriptome evolution, for each orthotype by performing Spearman rank-correlations on cell-type averaged (pseudobulk) transcriptome measurements. One-to-one orthologs across five rodent species (mouse, rat, deer mouse, hamster, and squirrel) that share all major DRG neuron orthotyeps were obtained from a mouse-referenced BioMart ortholog table, retaining only genes annotated as one-to-one orthologs in every species and yielding ∼12,984 unique orthologous gene sets. For each single-nucleus RNA-seq replicate, gene identifiers were converted to their mouse reference symbols via case-insensitive matching, datasets were subset to the ∼10,060 genes shared across all objects, and counts were pseudobulked by summing raw counts within each replicate × orthotype before normalizing to log-CPM. To construct transcriptome-based trees for individual orthotypes, the corresponding pseudobulk columns were variance-filtered, z-scored across samples, and converted to a distance matrix of 1 – Spearman’s rho (ρ) between samples, from which neighbor-joining trees were built; branch support was estimated by gene-level bootstrapping (1,000 resamples of genes with replacement), with clade support computed as the proportion of bootstrap trees recovering each clade. To quantify inter-orthotype transcriptome divergence, replicates were collapsed by species (summing raw counts within each species × cell type, then re-normalizing to log-CPM), and for each orthotype a cross-species divergence metric was computed as the mean pairwise distance (1 – ρ) across the five species. Uncertainty in this metric was assessed by gene bootstrapping (1,000 resamples), and per-orthotype divergence distributions were summarized as bootstrap quantile boxplots. C-Heat (Calca–) neurons were excluded from subsequent inter-orthotype analyses due to their extremely low cell numbers (fig. S3).

To measure inter-orthotype variability in transcriptomic divergence across species, we summed total tree branch lengths for each orthotype. These values represent quantitative measures of pseudobulk transcriptome distance or similarity that can be plotted on a tree to display pseudobulk transcriptome measures of relatedness. Notably, these trees did not necessarily recapitulate known species phylogeny (fig. S11B), rather suggesting substantial interspecies dynamicity in gene expression such that each species may adopt orthotype-characteristic transcriptomic profiles. This result suggests that some orthotypes have undergone significant transcriptomic divergence between some mammalian species relative to others. By contrast, the phylogenetic recapitulation exhibited by this analysis performed on spermatogonial cells suggests that distinct cell lineages may differ with regards to whether transcriptome distances recapitulate organismal phylogeny or not (78).

### Single-nucleus ATAC-seq Analysis

Single-nucleus ATAC-seq data were processed in R using Seurat (108) and Signac (113) packages.

#### Single-nucleus ATAC-sequencing UMAP embeddings

To generate single-nucleus ATAC-seq UMAP embeddings for each of several rodent samples, the Cell Ranger ARC peak-by-cell matrix and fragment file were loaded into a Signac chromatin assay annotated with Ensembl gene models, filtered to previously identified neuronal barcodes with transferred cell-type labels, and quality-filtered on ATAC counts, nucleosome signal, and TSS enrichment. Accessibility was then normalized by TF-IDF, reduced by latent semantic indexing (SVD on the top features), and embedded and clustered using the LSI dimensions (UMAP and graph-based Leiden clustering), after which non-neuronal or low-quality clusters were removed and the normalization, dimensionality-reduction, and clustering steps were repeated on the retained cells before UMAP embeddings were plotted.

#### ChromVAR analysis

For each of five rodent species (mouse, rat, deer mouse, hamster, and ground squirrel) that share all major DRG neuron orthotyeps, neuronal nuclei passing an initial fragment threshold (>500 ATAC fragments) were retained based on barcodes from the matched single-cell RNA-seq objects, fragments were quantified against the per-sample Cell Ranger ARC peak set (after removing peaks <20 bp or >10 kb), and cell-type labels were transferred from the RNA objects. Peaks were then re-called for each cell type using MACS3 and quantified into a new chromatin assay; for several species, peak coordinates were remapped from RefSeq accessions to the chromosome/scaffold names of the corresponding BSgenome and filtered to sequences within genome bounds, with mouse peaks additionally restricted to standard chromosomes and depleted of ENCODE blacklist regions. Per-base sequence statistics and JASPAR2020 vertebrate CORE motif matches were added to each object, and per-cell transcription factor (TF) motif accessibility was estimated using chromVAR. To compare motif activity across species, chromVAR deviation scores were collapsed into TF families (by normalizing TF names and stripping numeric and letter suffixes), family-level scores were averaged per cell type, and families were ranked by their variance across cell types in the mouse dataset to select the ten most variable families. For visualization, family-averaged chromVAR scores were min–max normalized within each sample and family across cell types and displayed as a dot plot faceted by TF family, with point color encoding normalized mean deviation and point size encoding the normalized fraction of cells with positive deviation, plotted for all five species side by side.

### Signac LinkedPeaks analysis

#### Mouse and rat sensor gene-peak linkage analysis

For each species, peak sets called by Cell Ranger ARC across samples were merged into a unified set of genomic ranges, and peaks shorter than 20 bp or longer than 10 kb were removed; fragments were then re-quantified against this common peak set in every sample. Nuclei were filtered to retain those with more than 500 ATAC fragments and to keep only neuronal barcodes previously identified in the corresponding single-nucleus RNA-seq objects, and the matched RNA and SCT assays together with their cell-type annotations were transferred onto the ATAC objects before all samples of a species were merged into a combined object. Peaks were then re-called per cell type on the merged data using MACS3, retaining only standard chromosomes and (for mouse) excluding ENCODE blacklist regions, and fragment counts were quantified against this refined peak set to build a new chromatin assay annotated with Ensembl gene models (mouse: EnsDb.Mmusculus.v79 with UCSC-style chromosome names; rat: EnsDb.Rnorvegicus.v105 with NCBI-style names). The resulting per-species objects, with cell types set as the active identities, were retained for downstream analyses.

For each species, peak GC content was computed and peak-to-gene links were identified for the sensor gene by correlating peak accessibility with SCT-normalized gene expression within a 500 kb window upstream and downstream of the gene (LinkPeaks, score cutoff 0.01). The resulting links and accessibility profiles were visualized per orthotype as coverage plots spanning the same flanking window. Linked peaks from one species were mapped to corresponding genomic loci/coordinates in the other species using the UCSC LiftOver online browser tool. The homologous coordinates were then replotted on coverage plots spanning the locus and 2 kb flanking windows.

### Histology: In Situ Hybridization (RNAscope) and Immunohistochemistry

Unless stated otherwise, mid-thoracic (T6 – T10, usually T8) DRGs were collected and used in studies contributing to Figs. 2 to 4 and figs. S4 to S10; lumbar level 4 (L4) DRGs were collected and used in studies contributing to Fig. 5 and fig. S12; and cervical level 7 (C7), mid-thoracic, and lumbar level 4 (L4) DRGs were collected and used in studies contributing to Fig. 6 and figs. S14 and S15.

#### In situ hybridization (ISH)

Individual DRGs from rodents were rapidly dissected out in DMEM-F12, HEPES (Thermo Fisher/Gibco, 11330-032; supplemented with 12.5 mM D-glucose and 1x pen/strep) on ice. Axial levels were identified using the T12 DRG as a landmark: the T12 DRG was defined as the ganglion immediately caudal to the last rib. Macaque thoracic (T8-T10) DRGs were obtained from the Oregon National Primate Research Center. Human thoracic (T8-T10) DRGs were obtained from AnaBios. DRGs were embedded in OCT (Fisher, 1437365) on ice, flash-frozen, and stored at −80 °C until sectioning on a cryostat. DRGs were sectioned at a thickness of 20 – 25 μm. Target transcripts were detected by RNAscope (Advanced Cell Diagnostics) using the manufacturer’s protocol. Briefly, after cryosectioning, slides were fixed in 4% prechilled PFA (paraformaldehyde; Sigma, P6148-500G, or EMS 32% aqueous solution, 15714-S) at 4°C for 15 mins, then dehydrated using serial concentrations of ethanol. Next the sections were digested using Protease Plus for 10 mins at RT. After washing, a mix of RNAscope probes was added on the slide and incubated at 40°C for 2 – 4 hours, followed by sequential AMP1, AMP2, and AMP3 steps, according to the protocol. Slides were mounted in DAPI Fluoromount-G (Southern Biotech, 0100-20) and imaged on a Zeiss LSM 900 confocal microscope.

The following RNAscope probes were used: Mm-Gfra2 (441481-C2), Mm-Rbfox3 (313311-C3), Mm-Th (317621-C3), Mm-Zfp521 (565831), Rn-Gfra2 (463041), Rn-Rbfox3 (436351-C3), Rn-Zfp521 (1299151-C2), Hg-GFRA2 (1299141-C2), Hg-RBFOX3 (541911-C3), Hg-ZNF521 (1299121-C1), Mfa-GFRA2 (466311), Mmu-RBFOX3 (870421-C3), Mmu-ZNF521 (1333631-C2), Hs-GFRA2 (463011), Hs-RBFOX3 (415591-C2), Hs-ZNF521 (565741-C3).

Mm-Trpa1 (400211-C3), Mm-Trpv1 (313331-C3), Mm-Cysltr2 (452621-C3), Mm-Etv1 (433281-C2), Mm-Gfra1 (431781), Mm-Piezo2 (400191-C3), Mm-Avil (498531), Mm-Mrgpra3 (548161-C3), Mm-Mrgprb4 (435781-C2), Rn-Trpa1 (312511-C3), Rn-Trpv1 (501161-C3), Rn-Cysltr2-O1 (1575841-C3), Rn-Etv1 (1209241-C1), Rn-Gfra1 (463031-C2), Rn-Piezo2 (549741-C3), Hs-TRPA1 (311951-C2), Hs-CYSLTR2 (543481-C2)

#### Cell-type proportion quantifications

For the data in Fig. 2 and fig. S4F, total numbers of neurons per DRG section were recorded by counting Rbfox3+/RBFOX3+ cells. Hair follicle density estimates from were derived from the literature (53, 54). For the data in fig. S15, total numbers of neurons per DRG section were recorded by counting Avil+ cells.

#### Sensor/receptor-cell type quantifications

To quantify sensor/receptor expression in distinct orthotypes using ISH data, we manually outlined individual DRG neuron cell bodies in Fiji (ImageJ; NIH, https://imagej.net/Fiji/Downloads) and measured the mean fluorescence level of all three channels (marker gene 1, marker gene 2, and sensor/receptor) in each of tens to hundreds of cells per section. Normalized signal from all channels was then quantified and three-dimensional results plotted using Prism (Prism 11, GraphPad, RRID:SCR_002798, https://www.graphpad.com/), with co-marker gene 1 on the x-axis, co-marker gene 2 on the y-axis, and the sensor/receptor fluorescence levels plotted by color. Individual marker gene fluorescence level distributions were examined and thresholds for demarcating cell types based on marker genes were determined. Finally, sensor/receptor fluorescence level distributions were examined and thresholds for defining a cell as sensor/receptor-positive versus -negative were determined.

#### Immunohistochemistry (IHC)

Animals were perfused with PBS followed by 4% PFA. Spinal columns were removed and DRGs and/or spinal cords dissected out. Back hairy skin was removed, treated with Nair hair removal lotion (NAIR hair removal cream, Church and Dwight Co., Princeton, NJ), and gently rubbed to remove hair and underlying fat. Skin was then washed and fixed in Zamboni fixative (picric acid-formaldehyde (PAF) fixative, Fisher, NC9335034) overnight (4°C). Tissue was embedded in OCT (Fisher, 1437365) or NEG-50, frozen and stored at –80°C. Tissue was cryosectioned and mounted on Superfrost plus slides (Fisher, 12-550-15). After drying at room temperature (RT) overnight, slides were rehydrated and washed with PBS, and blocked by 5% Normal Donkey Serum (Jackson, 017-000-121) for 1 – 2 hours at RT. Then primary antibody mix was added overnight at 4°C. Following incubation with primary antibody, slides were washed with 0.1% PBS-Triton, the secondary antibody mixed was added for 2 hours at RT. Finally, slides were washed in PBS, mounted in DAPI Fluoromount-G (Southern Biotech, 0100-20), and imaged using a confocal microscope (Zeiss LSM 900).

Primary antibodies used were goat anti-mCherry (Origene, AB0040-200; RRID:AB_2333093; 1:500), goat anti-GFP (US Biological Life Sciences, G8965-01E; 1:500), rabbit anti-TH (Sigma, AB152; RRID:AB_390204; 1:500), and rabbit anti-CGRP (Immunostar, 24112; RRID:AB_572217; 1:500). IB4-fluorophore conjugates (IB4, Alexa Fluor 647 conjugated; Thermo Fisher, I32450) were used at 1:500. Secondary antibodies included Donkey anti-goat (Alexa Fluor 488) (Thermo Fisher, A-11055; RRID:AB_2534102), Donkey anti-goat (Alexa Fluor 546) (Thermo Fisher, A-11056; RRID:AB_2534103), Donkey anti-rabbit (Alexa Fluor 488) (Thermo Fisher, A-21206; RRID:AB_2535792), and Donkey anti-rabbit (Alexa Fluor 546) (Thermo Fisher, A-10040; RRID:AB_2534016).

#### In situ hybridization-immunohistochemistry (ISH-IHC)

To accommodate joint in situ hybridization and immunohistochemistry on the same tissue sections, we performed the ISH protocol described above followed by the IHC protocol described above.

### Enhancer AAV (eAAV) Screening

#### Single-nucleus multiome ATAC-seq dataset processing and cell-type-specific enhancer nomination (PIASO pipeline)

Mouse DRG single-nucleus multiome ATAC-seq data were processed with PIASO (*114*) (https://github.com/genecell/PIASO). ATAC fragments were imported into a Cytome database file using piaso.pp.importFragments with per-nucleus transcription start site (TSS) enrichment computed inline against the mm10 TSS annotation; nuclei were retained if they had ≥ 1,000 fragments and a TSS enrichment ≥ 3, yielding 19,721 nuclei. Groups for peak calling were defined by two-level local Leiden clustering: a coarse partition (piaso.tl.leiden, resolution 0.25) followed by within-group refinement (piaso.tl.leiden_local, resolution 0.75), the latter re-embedding each coarse group in its own term frequency–inverse document frequency (TF-IDF) normalization (piaso.tl.run_TFIDF) followed by randomized singular value decomposition (SVD; 30 components). Peak calling was then performed by PICCO (piaso.tl.picco), operating directly on the compressed fragment store in Cytome. Peaks were quantified into a cell × peak count matrix (piaso.pp.quantifyPeakActivity) and TF-IDF-normalized. Cell-type labels were transferred from the independent analysis of the paired snRNA-seq multiome data, matched by cell barcode. Cell type-specific enhancers were nominated with COSG (*115*) applied to the TF-IDF peak layer (cosg.run_cosg_cytome), grouped by the RNA-derived cell-type annotation (mu = 100; expressed_pct = 0.05). From the ranked enhancer-candidate lists produced by COSG, we manually inspected the per-cell-type peak distributions in the corresponding bigWig tracks and selected the genomic regions surrounding the peak summits. To further prioritize candidates, we incorporated vertebrate sequence conservation, computed with phastCons (*116*, *117*) from the 60-way vertebrate alignment (mm10.60way.phastCons.bw, downloaded from http://hgdownload.cse.ucsc.edu/goldenpath/mm10/phastCons60way/).

#### AAV production and neonatal IP injection

AAV genome plasmids were constructed using standard cloning and molecular biology techniques (constructs generated in this paper: pAAV-mBG-mScarlet; pAAV-mBG-ReaChRmCitrine; pAAV-TRE3g-GCaMP8s-WPRE-bGHpA). AAVs (serotype PHP.S) were packaged and concentrated using transient transfection of PHP.S, pHelper, and AAV-genome plasmids into HEK 293T cells. Supernatants were collected at 72 and 120 hours post-transfection. At 120 hours post-transfection, cells were scraped off plates and collected. 293T cell pellets were extracted using a lysis buffer containing salt active nuclease (ArcticZymes) in 40 mM Tris, 500 mM NaCl and 2 mM MgCl_2_ pH 8 (SAN buffer). Viral supernatants were concentrated via 8% PEG/500 mM NaCl precipitation and re-suspended in SAN buffer. Cleared viral lysates were then loaded onto an iodixanol gradient (Optiprep; OptiPrep™ Density Gradient Medium, Sigma-Aldrich, D1556-250ML), centrifuged, and concentrated using Amicon filters with a 100-kD molecular cutoff to a final volume of approximately 15 – 20 µL. AAVs were injected intraperitoneally into pups at postnatal days 0 – 2.

#### eAAV injections

For eAAV screening, 5e11 – 2e12 genome copies (GC) per eAAV were injected into mouse pups. For eAAV-mediated behavioral testing, 2e11 – 4e12 GC eAAV(PHP.S)-C-LTMRe1L-ReaChR-mCitrine and eAAV(PHP.S)-C-LTMRe1L-mScarlet was injected into mouse pups, and 2e12 – 4e13 GC eAAV(PHP.S)-C-LTMRe1L-ReaChR-mCitrine and eAAV(PHP.S)-C-LTMRe1L-mScarlet was injected into rat pups. For eAAV-mediated Ca2+ imaging physiology, 5e11 – 1e12 GC eAAV(PHP.S)-C-M/H(Mrgprd)e7-tTA was injected along with 1e12 GC AAV(PHP.S)-TRE-GCaMP8s into mouse pups, while 1e12 – 2e12 GC eAAV(PHP.S)-C-M/H(Mrgprd)e7-tTA was injected along with 1e12 – 2e12 GC AAV(PHP.S)-TRE-GCaMP8s into rat pups.

### Behavioral Analyses

#### Animals

Mice and rats were maintained on a standard 12-hour light/dark cycle (lights on 7:00 AM, lights off 7:00 PM) with ad libitum access to food and water. Before optogenetic stimulation, animals were lightly shaved from behind the ears to just past the shoulders. Testing was conducted in a custom behavioral chamber consisting of a 10 × 5 inch enclosure with three black matte acrylic walls and an optically clear front panel, with a clear acrylic center divider allowing visual contact between animals on either side. The floor was white matte acrylic covered with a disposable soft cloth surface that was changed between each cage to prevent olfactory contamination and maintain consistent testing conditions. The black matte walls minimize reflections; the optically clear front panel allows observation and recording during testing without obstruction. Prior to behavioral testing, animals underwent a structured habituation protocol ≥ 2 days. Each session consisted of 90 minutes of room habituation followed by 30 minutes of chamber habituation, after which animals were returned to their home cage.

#### Optogenetic stimulation, testing protocol, and behavioral scoring

Transcutaneous optogenetic stimulation was delivered using a LEDD1B LED driver with an M470F3 470nm LED coupled to an M41L01 patch cable (600 µm core diameter, 0.48 NA, Thorlabs). The fiber tip was held at a fixed distance of six inches from the animal’s skin surface, producing a spot area of ∼0.71 cmZ. Tip power was measured at each nominal setting using a calibrated power meter prior to each experiment. For mice, stimulation was delivered at powers of 1.5–3.5 mW (2.31–4.97 mW/cmZ). The 1.5 mW setting (2.31 mW/cmZ) served as a sub-threshold light-on control producing no behavioral response. For rats, stimulation was delivered at 4–12 mW (6.38–17.65 mW/cmZ). Irradiance values were calculated from measured tip power and a fixed 0.71 cmZ spot area at 6-inch tip-to-animal distance.

For each session, mice and rats were allowed a brief settling period of around five minutes, then the LED fiber tip was positioned above the neck and shoulder region of the animal. Light was delivered in brief movements across the stimulation area while the experimenter monitored for wet dog shake (WDS) responses in real time. Sessions were recorded from front and top camera views using DMK 33UX287 cameras (The Imaging Source) at 120 frames per second in uncompressed format.

Videos were scored manually frame by frame using VirtualDub software. For each animal and stimulation trial, several parameters were recorded, including wet dog shake response (positive/negative), stimulus onset frame or time, event onset (head cock, full WDS, end) frame or time, latency to WDS onset (in seconds). A WDS was defined as a discrete bout of rapid oscillatory movements of the head and upper trunk, consistent with previously published criteria (50, 59).

#### Oil/water stimulation (rats)

Animals were habituated in the acrylic chambers as above and settled for at least five minutes before stimulation. Once animals appeared at ease, room temperature water or sunflower seed oil (Sigma, S5007) was applied onto the upper back of the animals, targeting the region between the shoulder blades. Water and oil stimuli were applied on different days and thus during separate recording sessions.

#### Oil/water stimulation (naked mole-rats)

Naked mole-rats were habituated for five minutes in a 5 × 2 × 6 inch (length × width × height) acrylic arena. Following habituation, either mineral oil (Sigma-Aldrich, M8410-1L) or water was applied to the upper back/nape of the animal’s neck. Animals were then recorded for an additional five minutes at 240 frames per second. The number of WDSs, duration of scratching bouts, and duration of grooming bouts during the five-minute post-application period were quantified from video recordings. Although naked mole-rats never performed WDSs following oil or water stimuli, each one always immediately attended to the oil and water stimuli right after stimulus presentation, often by orienting their head in the direction from which the stimulus was applied, indicating that they sensed the oil or water applied. Naked mole-rats spent most of each behavioral recording session displaying typical locomotion, burrowing, and digging behaviors.

### *In Vivo* Epifluorescence Ca^2+^ Imaging

Mice of both sexes at 4 – 6 weeks were anesthetized with inhalational isoflurane (Covetrus, 29405) throughout the experiments. Back hair was shaved, back skin disinfected, leg hair removed, and an incision was made over the lumbar vertebrae. Paravertebral muscles along the L4 vertebral spine were dissected, then the bone on top of the L4 DRG was removed by rongeur. A spinal clamp was used to stabilize the spinal column. The surgical preparation was then placed under an upright epifluorescence microscope (Zeiss Axio Examiner) with a 10X air objective (Zeiss Epiplan, NA=0.20). The light source was a 470nm LED (M470L5, Thorlabs) with an LED driver (LEDD1B, Thorlabs). A CMOS Camera (CS505MU1, Thorlabs) was triggered at five or ten frames per second with a 50 ms exposure time. All the recorded stimuli were synchronized with the camera and LED using a DAQ board (National Instrument, NI USB-6343).

#### Stimuli

Chemical stimuli were prepared by dissolving in 0.9% saline (0.9% Saline Solution, Teknova, S5825) shortly before application. Stock capsaicin (Sigma-Aldrich, M2028-250MG) was dissolved in DMSO at 10% weight/volume. Stock capsaicin and AITC (allyl isothiocyanate, Sigma, 377430-500G) were diluted in 0.9% saline and applied onto the DRG in succession from 1:10,000, then 1:1,000, then 1:100, then 1:10, until Ca^2+^ responses were observed. Given the volatility of AITC and imperfect solubility of capsaicin in saline, we predicted that different concentrations could elicit Ca^2+^ responses across trials, even within the same animal (e.g., between the left and right L4 DRGs). In the case of all mouse DRGs, responses to capsaicin were not observed at any concentrations, including at 1:10, while responses to AITC were readily observed at concentrations usually orders of magnitude lower. In the case of rat DRGs, clear responses to both AITC and capsaicin were typically easily observed at concentrations orders of magnitude lower than 1:10. The first concentration at which responses were observed constituted the last application of that chemical stimulus to avoid over-stimulating DRG neurons and triggering cytotoxicity. Between application of different chemical stimuli, three washes of 20 µL saline were applied, and DRGs were allowed to recover for ten minutes with no stimuli. Cotton swabs were used to gently remove excess chemical or saline stimuli between trials or during washes.

All DRGs were stimulated with a standard sequence: mechanical stimulation (leg pinch) first, followed by saline (intended as a chemical negative control), followed by AITC, then capsaicin. AITC and capsaicin application order was reversed in some animals, with no obvious differences. After all chemical stimulations were concluded, a final round of mechanical (pinch) stimulation was often performed to confirm the health of the DRG neurons at the end of the trial. During all trials, stimuli were not applied during frames 1 – 100, allowing these frames to constitute a baseline period for each stimulus trial. During mechanical stimulation, forceps were used to pinch the hairy hindlimb during frames 101 – 300 and the hindpaw during frames 301 – 500. Videos were captured at 10 Hz during mechanical stimulus trials and 5 Hz during chemical stimulus trials. All chemical (including negative control saline) stimuli were applied onto the DRG between frames 101 and 150 and liquid was removed from the DRG by frame 200. Chemical responses were quantified starting from frame 201. Mechanical stimulus trials lasted 60 seconds, with the last 10 seconds not having any stimuli; mechanical stimulus trials thus yielded 500 frames in total. Chemical stimulus trials lasted 120 seconds, with the second 20 seconds (frames 101 – 200) being spliced out due to chemical application and removal; chemical stimulus trials thus also yielded 500 frames in total.

#### Calcium imaging processing and analysis

For imaging analysis, initial processing was conducted in Fiji (ImageJ; NIH, https://imagej.net/Fiji/Downloads). Motion correction and spatial high-pass filtering was conducted using a custom-written macro utilizing ImageJ plugins moco and Unsharp Mask. Regions of interest corresponding to cells that exhibited baseline signal and/or calcium response to any stimuli were selected, their coordinates aligned across videos, and their mean fluorescence intensity measurements recorded from across all stimulus trials.

The mean fluorescence intensity measurements from Fiji were further processed and analyzed using MATLAB. To calculate ΔF/F, F was defined for each trial as the minimum of the mean or median of the fluorescence intensity during the baseline 100 frame period before each stimulus. ΔF/F measurements were then normalized for plotting in heatmaps. For the heatmaps in Figs. 5 and fig. S13, data were filtered to include only cells that responded to at least one chemical and then ordered by responses to AITC then to capsaicin. For the heatmap in Fig. 6, data were filtered to include only cells that responded to at least one chemical and to mechanical stimulation, grouped by responsiveness to mechanical stimulation in the 100 – 300 frame range (the hairy hindlimb mechanical stimulation period), then the 300 – 500 frame range (the hindpaw mechanical stimulation period), and then ordered by responses to AITC then to capsaicin.

---

**Fig. S1.**
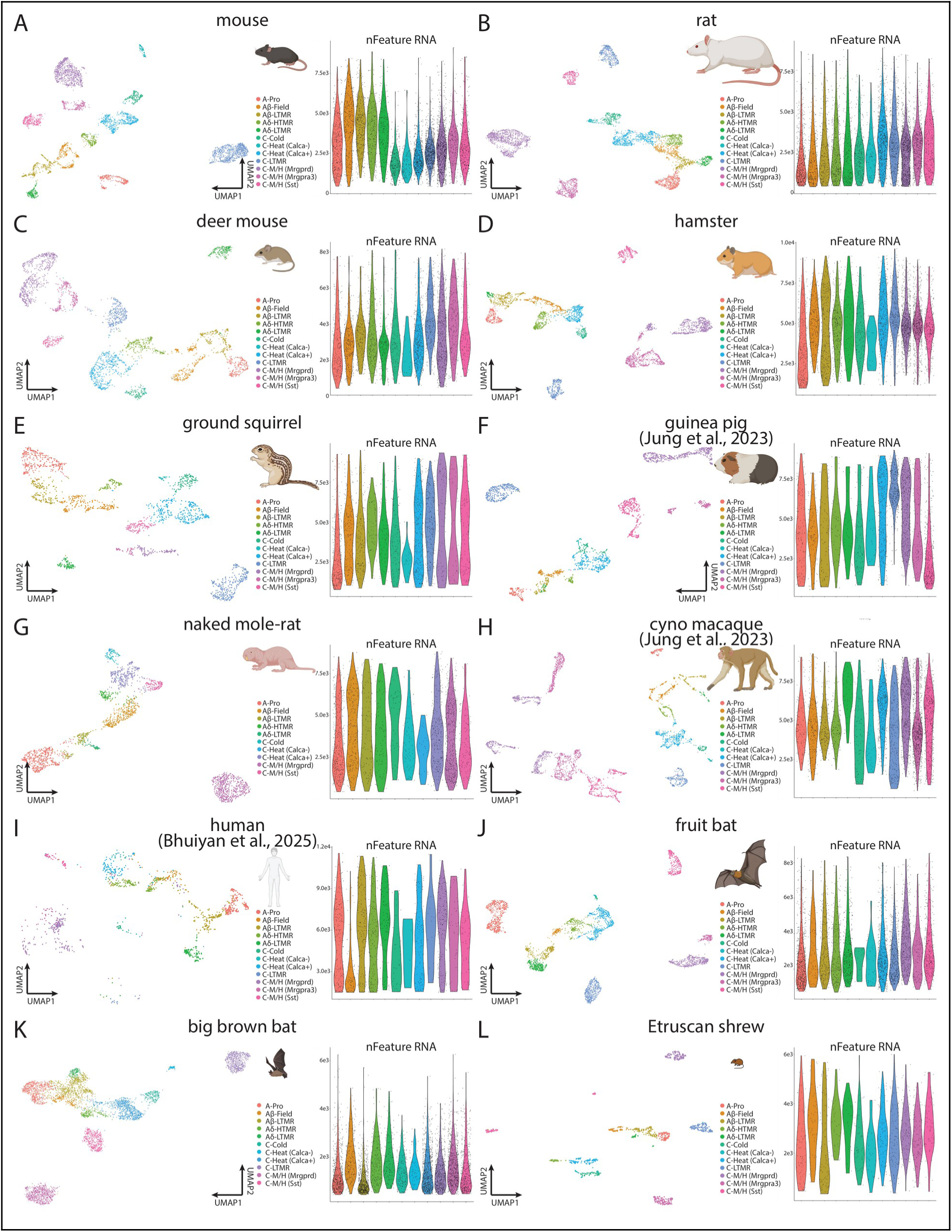
Example DRG neuron single-nucleus RNA-sequencing replicate datasets. (**A** to **L**) Left, Unform Manifold Approximation and Projection (UMAP) plots of DRG neuron single-nucleus RNA-sequencing datasets. Datasets plotted are from single biological replicates from distinct individual animals for all species except guinea pig (F), naked mole-rat (G), cynomolgous macaque (H), and human (I). Guinea pig and cynomolgous macaque datasets are from (*33*) and the human dataset is from (*35*). UMAP datapoints are colored by DRG neuron orthotype. Right, quality control nFeature RNA (genes per cell) metric plotted for each orthotype of each replicate for each species.

**Fig. S2.**
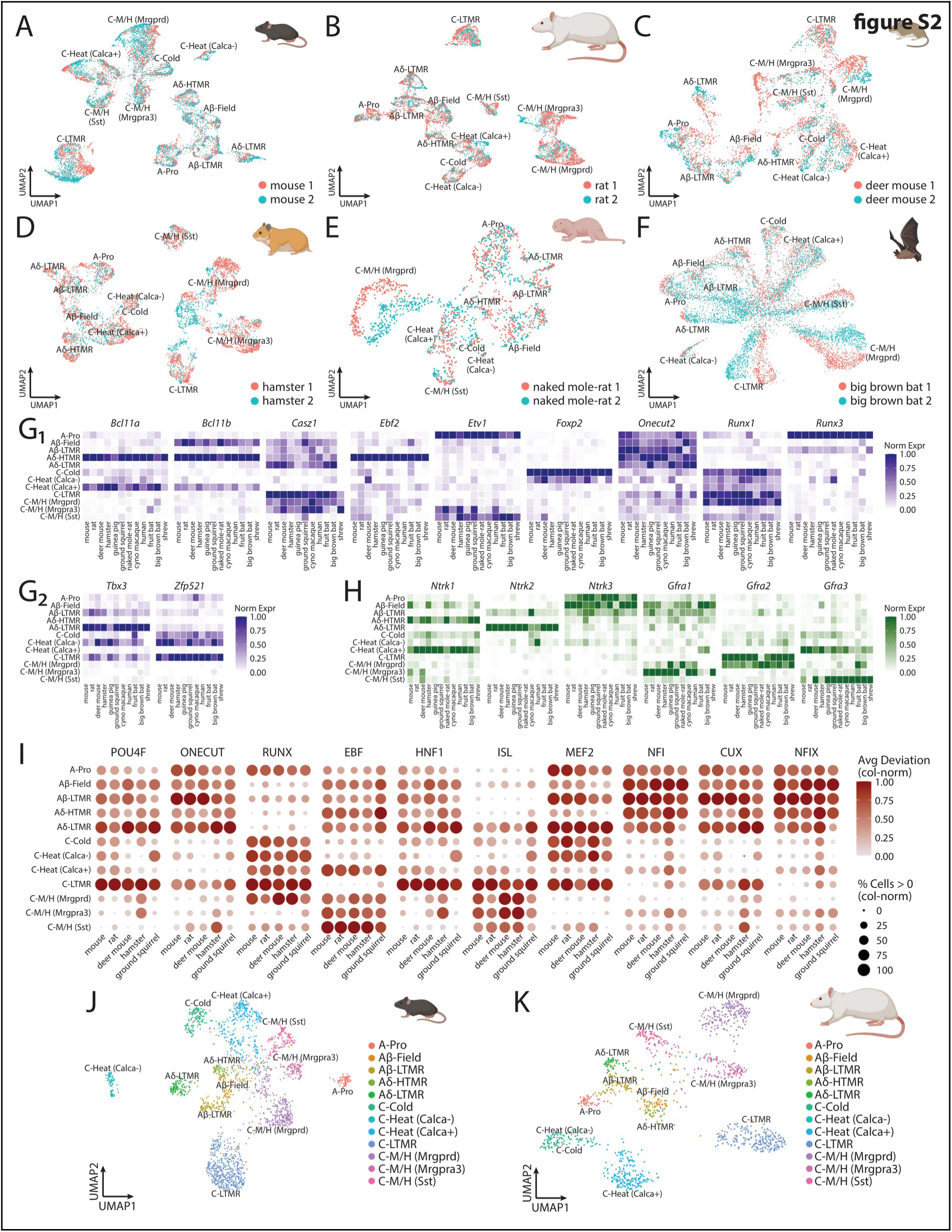
Additional evidence of DRG neuron-type conservation across species from single-nucleus RNA- and ATAC-sequencing datasets. (**A** to **F**) Comparison of biological replicates from different individual animals via UMAP plots of gene expression (RNA) datasets following within-species dataset merging (Seurat *merge* function), rather than integration, to preserve and display, rather than hide, any batch effects and technical variation. UMAP plot datapoints are colored by replicates (blue, red) and DRG neuron orthotypes are labeled on each plot. Sample species include mouse (A), rat (B), deer mouse (C), hamster (D), naked mole-rat (E), and big brown bat (F). Overlap between distinct replicates within clusters suggests biological similarity and high and consistent data quality across replicates. (**G** and **H**) Orthotype-specific expression patterns of transcription factor (G, purple) and growth factor receptor (H, green) genes across species. Gray circles, missing cell types (see Fig. 2 and fig. S3). (**I** and **J**) UMAP plots of ATAC datasets from example mouse (I) and rat (J) multiome datasets. Datapoints are colored by cell type.

**Fig. S3.**
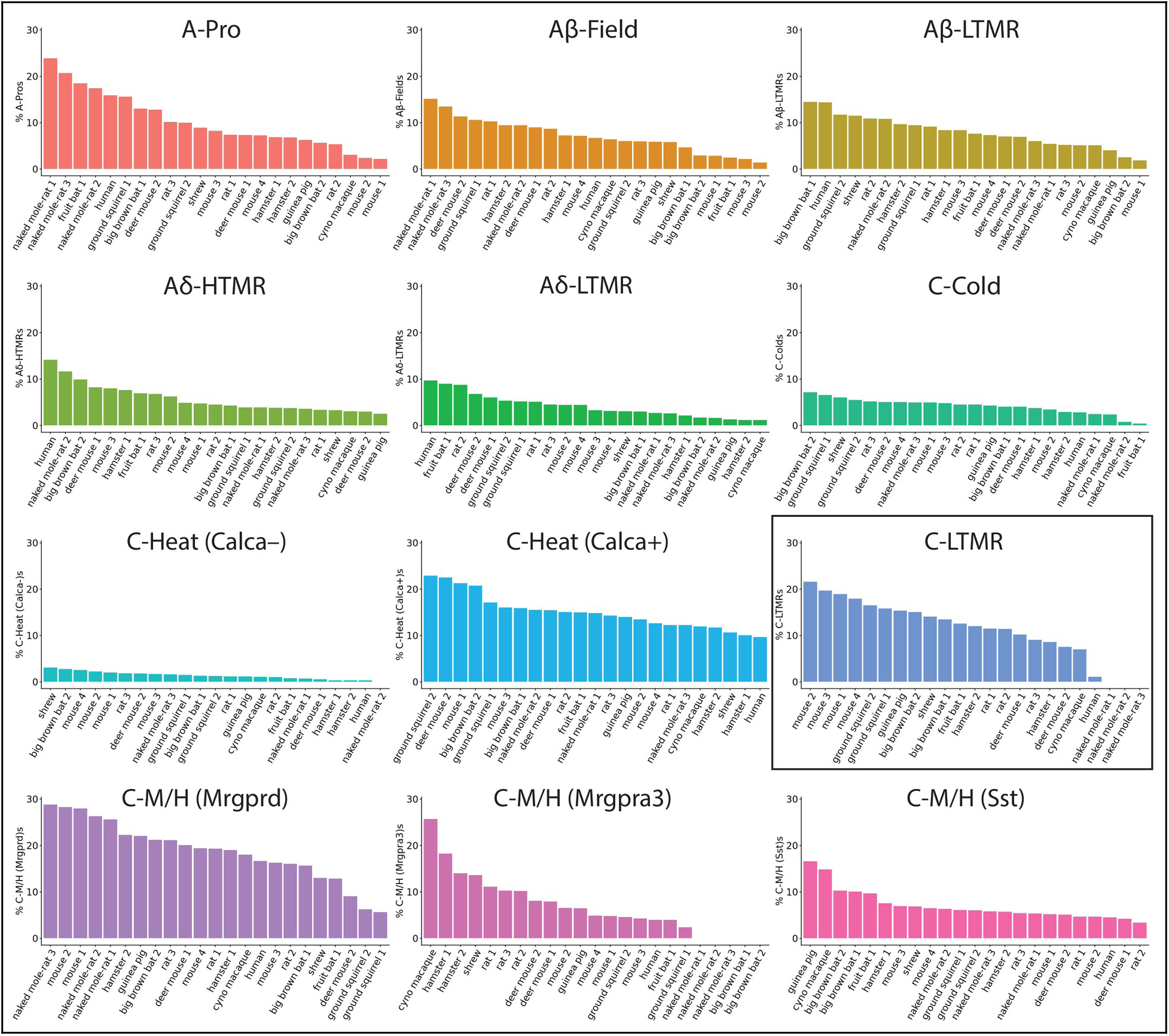
DRG neuron orthotype proportions of individual samples across species. DRG neuron orthotype proportions of individual sample replicates across species. Individual replicates are ordered by cell-type proportion within that replicate.

**Fig. S4.**
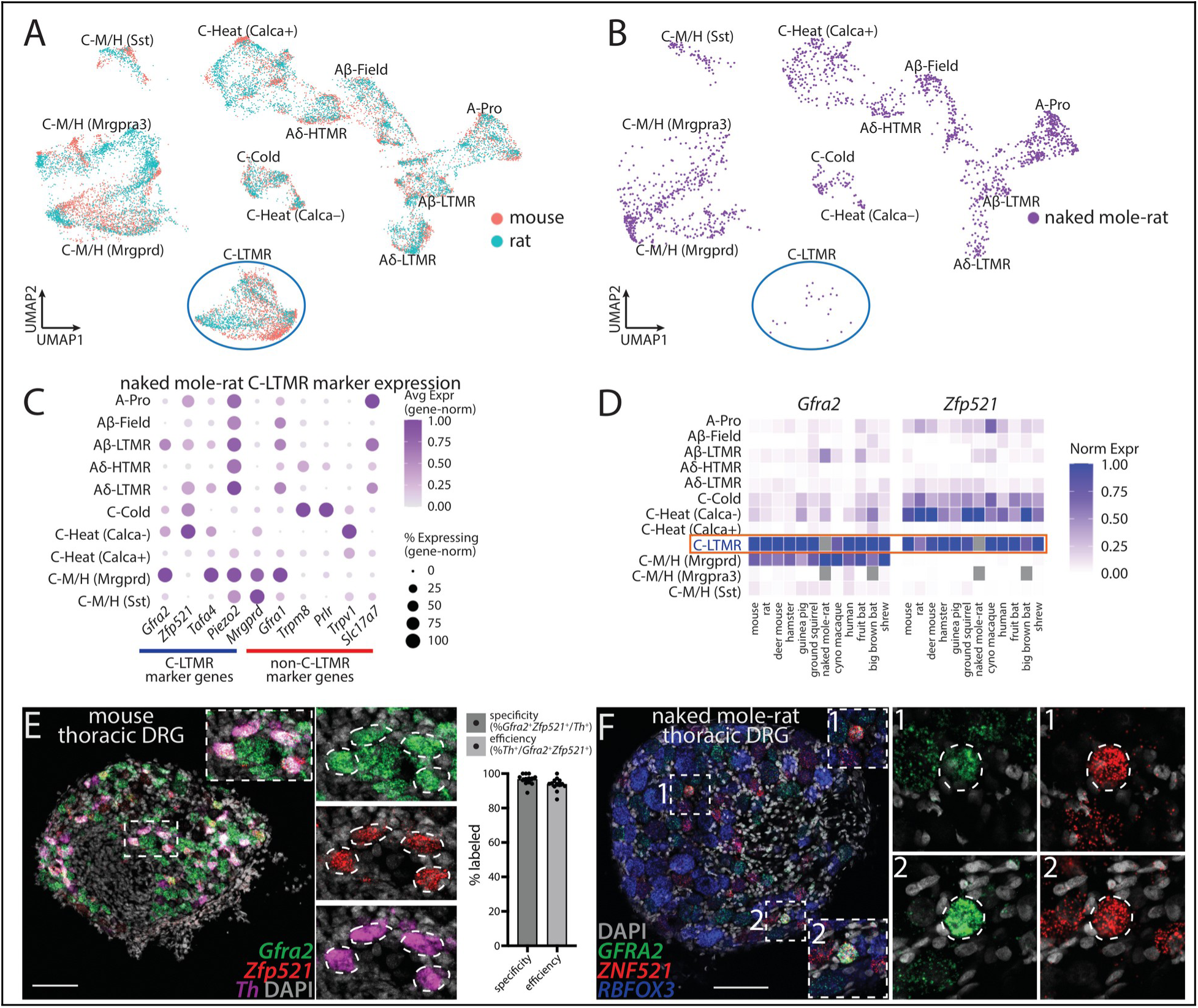
Additional characterization of C-LTMR neuron proportions across species. (A) Mouse (red) and rat (blue) samples in the pan-species UMAP embedding (see Fig. 1B). Note abundance of neurons from both mouse and rat in the C-LTMR cluster (circled in blue). (B) Naked mole-rat samples in the pan-species UMAP embedding (see Fig. 1B). Note relative paucity of neurons in the C-LTMR cluster (circled in blue). (C) Dot plot of C-LTMR and other DRG neuron orthotype marker genes in naked mole-rat samples. (D) *Gfra2*/*GFRA2* and *Zfp521*/*ZNF521* expression patterns across DRG neuron orthotypes and species shows that these two genes are conserved co-markers of C-LTMRs (orange box) across species. Gray boxes, missing cell types. (E) *In situ* hybridizations of mouse mid-thoracic DRGs showing overlap between *Gfra2* (green) and *Zfp521* (red), co-markers of C-LTMRs, and *Th* (magenta), a canonical marker of C-LTMRs in mice (*24*). Left, confocal images of *in situ* hybridizations. Right, quantifications; mean ± SEM. (F) *In situ* hybridization of naked mole-rat mid-thoracic DRGs against *GFRA2* (green) and *ZNF521* (red), co-markers of C-LTMRs, along with pan-neuronal marker *Rbfox3*/*RBFOX3* (NeuN, blue), sometimes reveal a small number (e.g., 1 – 5) of *GFRA2*^+^/*ZNF521*^+^ putative C-LTMRs per DRG, as outlined in dotted white lines in insets (right). Scale bars, 100 µm.

**Fig. S5.**
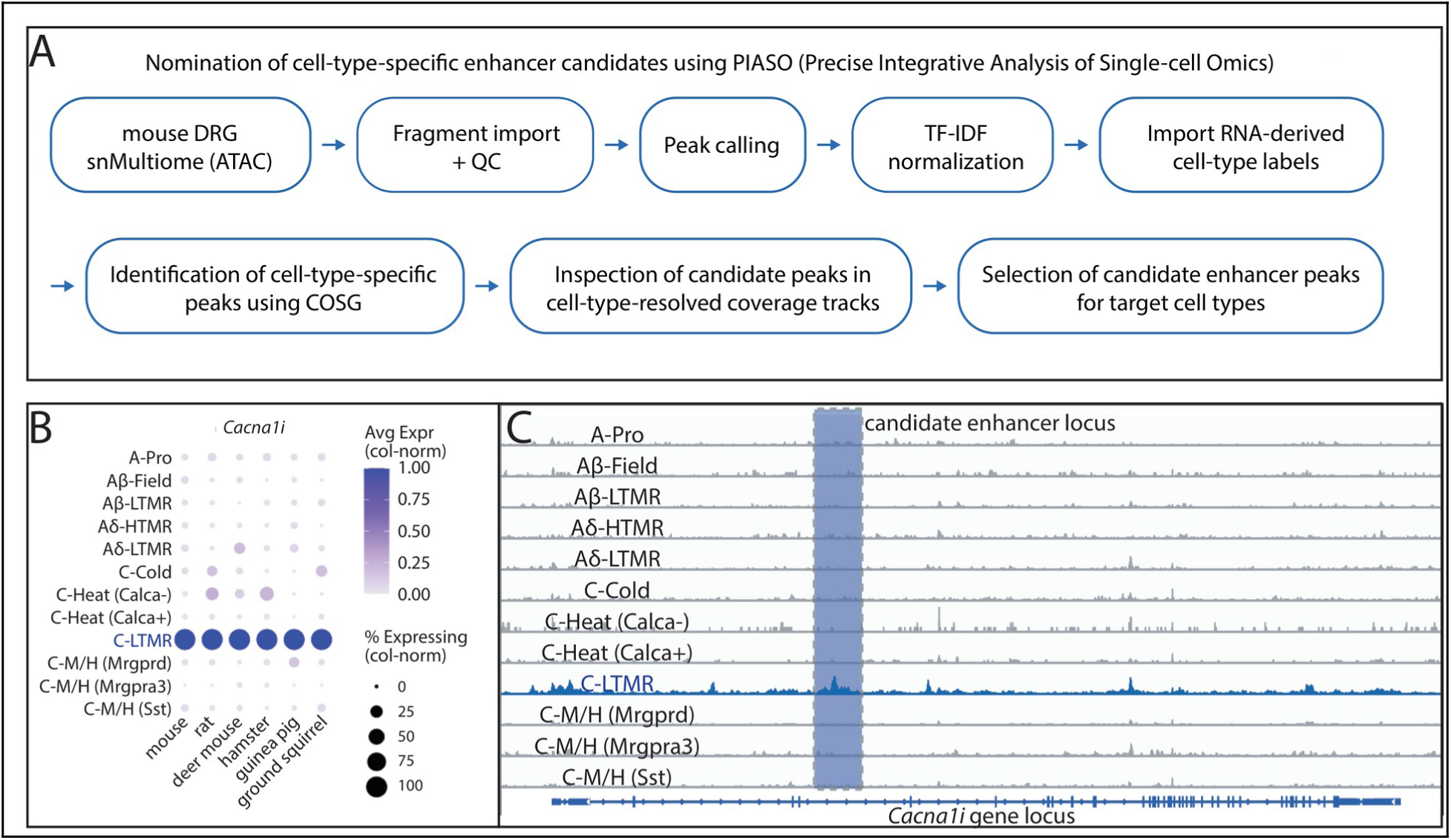
Enhancer candidate selection pipeline for C-LTMR neuron targeting. (A) Schematic illustrating PIASO pipeline for candidate enhancer selection. ATAC fragments were imported, with quality control per nucleus by fragment count and TSS enrichment; accessible peaks were then called. A cell × peak matrix was assembled and normalized by TF-IDF. Cell-type annotations transferred from the paired snRNA-seq analysis (matched by cell barcode) were used to group nuclei, and cell-type-specific marker peaks were identified with COSG. Candidate peaks were then inspected in cell-type-resolved coverage tracks, and enhancer candidates for each target cell type were selected from the regions surrounding peak summits. Vertebrate phastCons conservation was used to further prioritize candidates. TF-IDF, term frequency–inverse document frequency. (B) Dot plot showing specific expression of C-LTMR marker gene *Cacna1i* across rodent species. (C) Genome tracks showing enrichment of ATAC signal in top C-LTMR candidate *cis*-regulatory element in an intron of C-LTMR marker gene *Cacna1i* relative to other DRG neuron types. Blue shaded region corresponds to selected candidate sequence. See also Fig. 3A.

**Fig. S6.**
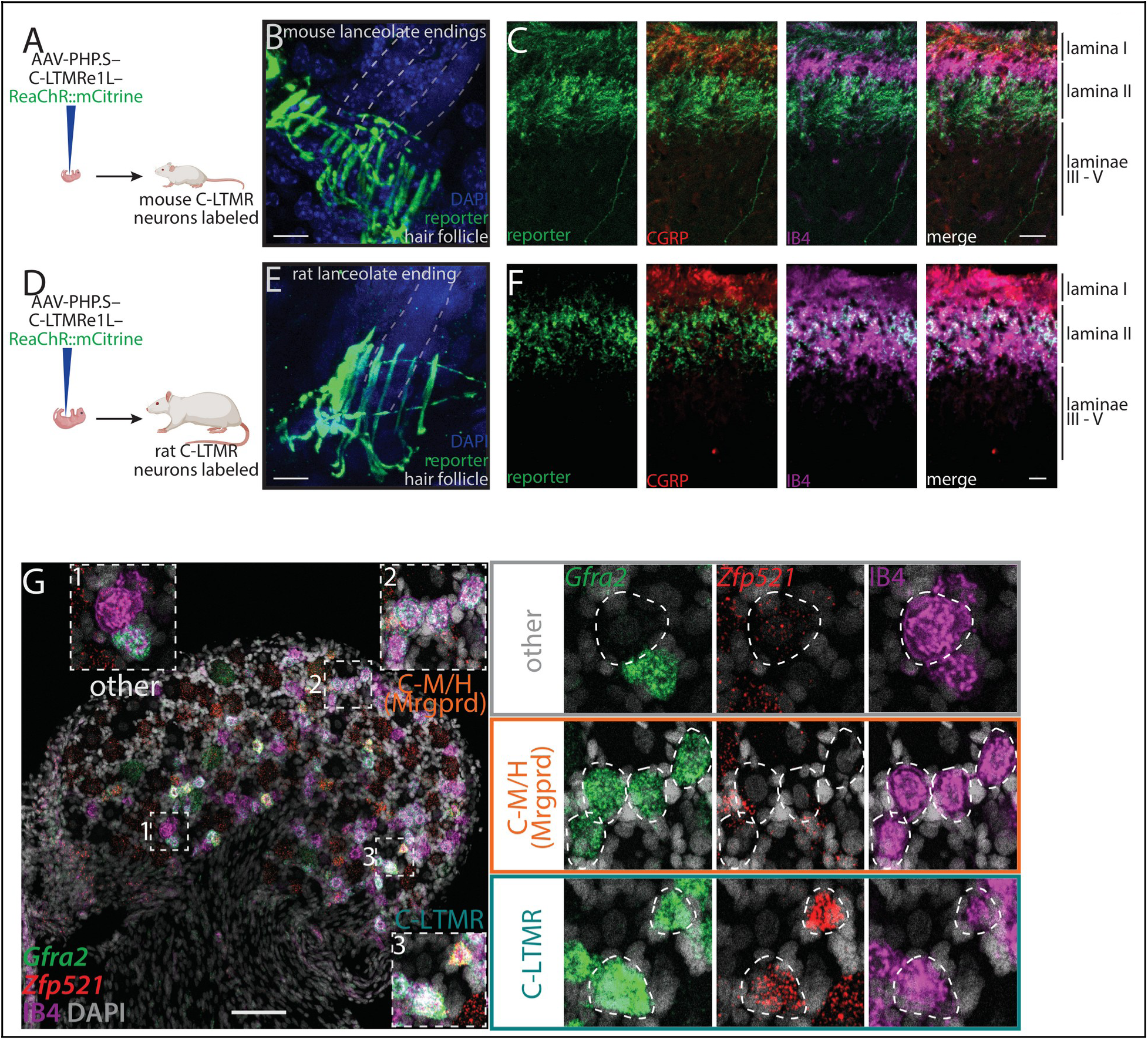
eAAV-mediated characterization of C-LTMR peripheral and central anatomy across mice and rats. (**A** to **F**) Schematics of eAAV-mediated C-LTMR targeting in mice (A) and rats (D). Labeling of lanceolate hair follicle endings in hairy skin of mice (B) and rats (E). eAAV-mediated C-LTMR neuron labeling in mice (C) and rats (F) reveals central terminals targeting spinal cord dorsal horn lamina II, ventral to lamina I marker CGRP (see also fig. S3D). Scale bars, 20 µm. (G) *In situ* hybridization characterization of IB4-binding rat DRG neurons. Rat IB4-binding neurons are a more diverse population than in mice. In mice, almost all (96%) IB4-binding neurons express *Mrgprd*, a marker of mouse C-M/H (Mrgprd) neurons (see fig. S12, B and C), while in rats, IB4-binding neurons (right insets) include *Gfra2*^+^/*Zfp521*^–^ putative C-M/H (Mrgprd) neurons, as well as *Gfra2*^+^/*Zfp521*^+^ putative C-LTMR neurons and other *Gfra2*^–^ neurons. Thus, IB4 binding may not mark the same DRG neurons and spinal laminae and sublaminae in rats as in mice (*118*, *119*). Scale bar, 100 µm.

**Fig. S7.**
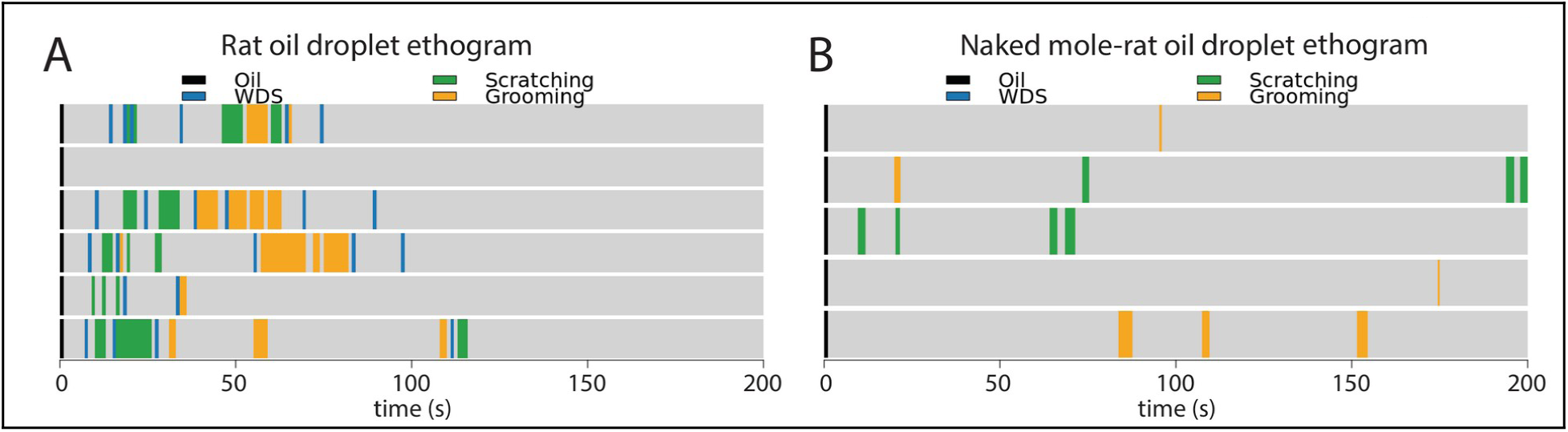
Characterization of rat and naked mole-rat somatomotor behavioral responses to oil application on the upper back. (**A** and **B**) Ethograms of rat (A) and naked mole-rat (B) responses to oil droplets targeted to the upper back. WDS, wet-dog shake. Black, blue, green, and yellow bars correspond to oil stimulus presentation, WDS bouts, bouts of scratching using the hindlimb, and bouts of facial and/or upper trunk grooming, respectively. Rats frequently perform WDS following oil stimulus presentation and other grooming/scratching behaviors, while naked mole-rats never perform WDS but do occasionally engage in other scratching and grooming behaviors.

**Fig. S8.**
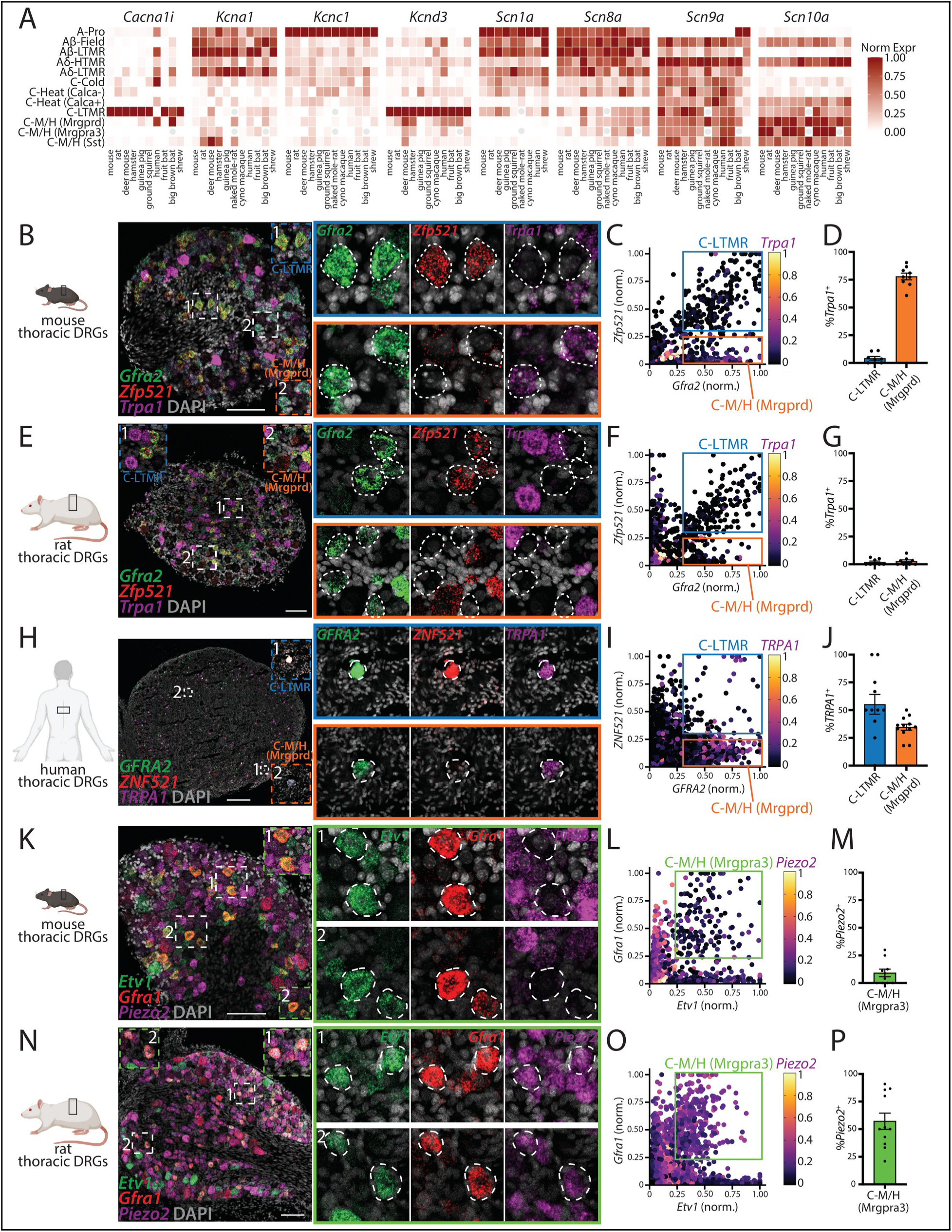
Additional validation of within-orthotype sensor shuffling across species. (A) Gene expression plot of orthotype ion channel expression across species. Gray circles, missing cell types (see Fig. 2 and fig. S3). (**B** to **J**) *In situ* hybridization validation of *Trpa1* expression patterns in C-LTMR (*Gfra2*^+^/*Zfp521*^+^) and C-M/H (Mrgprd) (*Gfra2*^+^/*Zfp521*^–^) neurons across mouse (B), rat (E), and human (H) mid-thoracic DRGs. (C), (F), and (I): Quantification of neuronal *Trpa1* expression (1001, 1540, and 4367 neurons from 9, 9, and 12 sections from 3, 4, and 4 DRGs from 2, 3, and 2 mice, rats, and humans, respectively). (D), (G), and (J): Percentages of *Trpa1*^+^ neurons (9, 9, and 12 DRG sections from mice, rats, and humans, respectively); mean ± SEM. (**K** to **P**) *In situ* hybridization validation of *Piezo2* expression patterns in C-M/H (Mrgpra3) (*Etv1*^+^/*Gfra1*^+^) neurons across mouse (K) and rat (N) mid-thoracic DRGs. (L) and (O): Quantification of neuronal *Piezo2* expression (1294 and 3070 neurons from 10 and 11 sections from 5 and 5 DRGs from 3 and 3 mice and rats, respectively). (M) and (P): Percentages of *Piezo2*^+^ neurons (10 and 11 DRG sections from mice and rats, respectively); mean ± SEM. Scale bars, 100 µm (B, E, K, N), 500 µm (H).

**Fig. S9.**
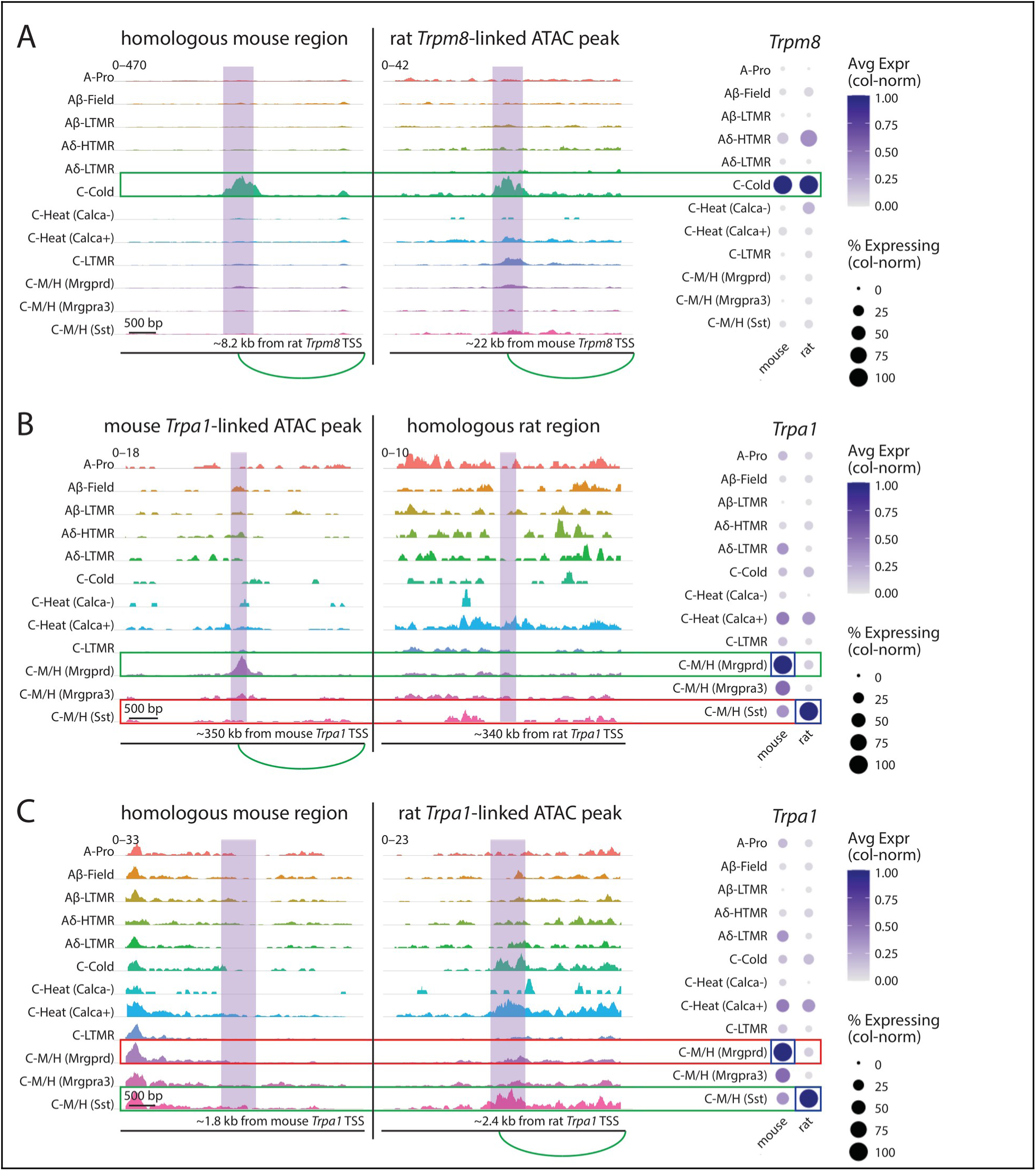
Species-specific utilization of genomic elements that may regulate sensor expression. (A) Left and middle, genome tracks showing ATAC signal across DRG neuron orthotypes in mouse (left) and rat (middle) genomic coordinates surrounding a rat *Trpm8*-linked ATAC peak (see Methods) and the corresponding homologous mouse genomic region. Right, dot plot showing *Trpm8* expression in mouse and rat DRG neuron orthotypes. C-Cold neurons, which express high *Trpm8* in both species, exhibit large, clear peaks near *Trpm8* in both rats and mice. (B) Left and middle, genome tracks showing ATAC signal across DRG neuron orthotypes in mouse (left) and rat (middle) genomic coordinates surrounding a mouse *Trpa1*-linked ATAC peak (see Methods) and the corresponding homologous rat genomic region. Right, dot plot showing *Trpa1* expression in mouse and rat DRG neuron orthotypes. Note that C-M/H (Mrgprd) neurons have the highest *Trpa1* expression in mice, while C-M/H (Sst) neurons have the highest *Trpa1* expression in rats. While mouse C-M/H (Mrgprd) neurons exhibit a clear peak linked to *Trpa1* expression, neither rat C-M/H (Mrgprd) neurons, which do not express appreciable *Trpa1*, nor rat C-M/H (Sst) neurons, which do express *Trpa1*, exhibit an ATAC peak at the homologous location. (C) Left and middle, genome tracks showing ATAC signal across DRG neuron orthotypes in mouse (left) and rat (middle) genomic coordinates surrounding a rat *Trpa1*-linked ATAC peak (see Methods) and the corresponding homologous mouse genomic region. Right, dot plot showing *Trpa1* expression in mouse and rat DRG neuron orthotypes. Note that C-M/H (Sst) neurons have the highest *Trpa1* expression in rats, while C-M/H (Mrgprd) neurons have the highest *Trpa1* expression in mice. While rat C-M/H (Sst) neurons exhibit a clear peak linked to *Trpa1* expression, neither mouse C-M/H (Mrgprd) nor C-M/H (Sst) neurons exhibit an ATAC peak at the homologous location.

**Fig. S10.**
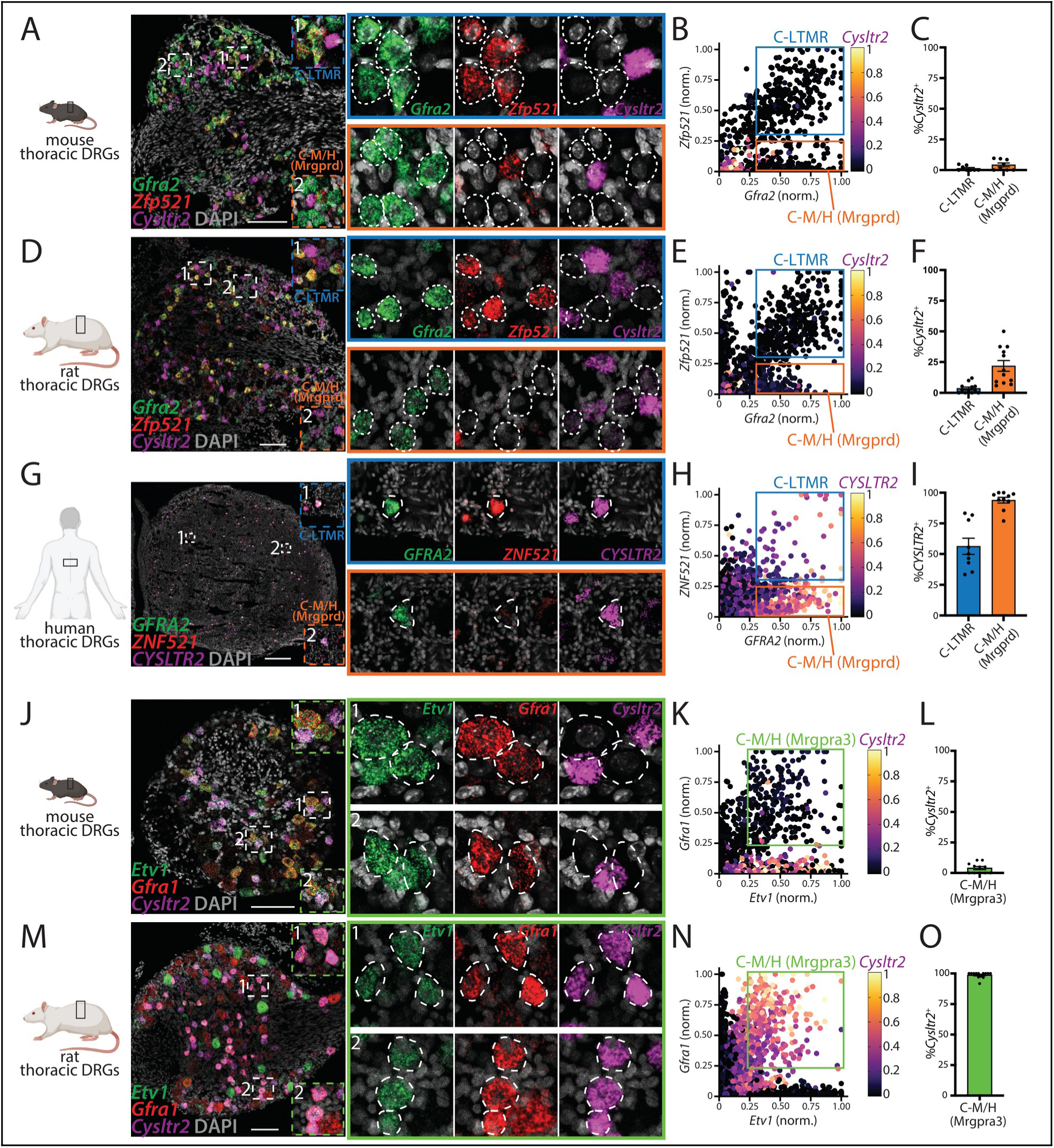
Validation of immune receptor shuffling across species. (**A** to **I**) *In situ* hybridization validation of *Cysltr2* expression patterns in C-LTMR (*Gfra2*^+^/*Zfp521*^+^) and C-M/H (Mrgprd) (*Gfra2*^+^/*Zfp521*^–^) neurons across mouse (A), rat (D), and human (G) mid-thoracic DRGs. (B), (E), and (H): Quantification of neuronal *Cysltr2* expression (1132, 2468, and 3430 neurons from 8, 12, and 10 sections from 4, 4, and 4 DRGs from 2, 2, and 2 mice, rats, and humans, respectively). (C), (F), and (I): Percentages of *Cysltr2*^+^ neurons (8, 12, and 10 DRG sections from mice, rats, and humans, respectively); mean ± SEM. (**J** to **O**) *In situ* hybridization validation of *Cysltr2* expression patterns in C-M/H (Mrgpra3) (*Etv1*^+^/*Gfra1*^+^) neurons across mouse (J) and rat (M) mid-thoracic DRGs. (K) and (N): Quantification of neuronal *Cysltr2* expression (1147 and 2488 neurons from 14 and 13 sections from 6 and 5 DRGs from 3 and 3 mice and rats, respectively). (L) and (O): Percentages of *Cysltr2*^+^ neurons (14 and 13 DRG sections from mice and rats, respectively); mean ± SEM. Scale bars, 100 µm (A, D, J, M), 500 µm (G).

**Fig. S11.**
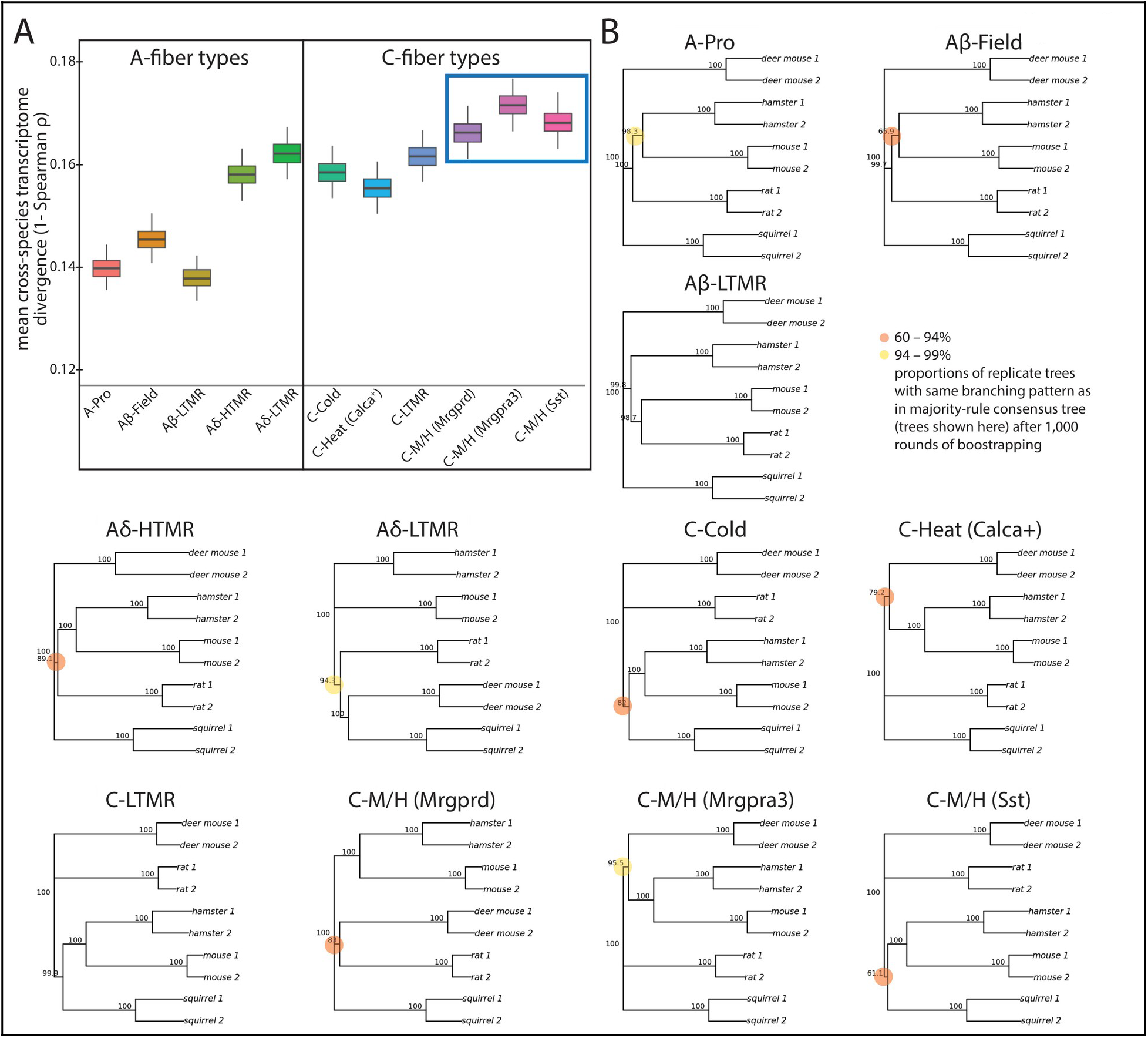
Characterization of DRG neuron orthotype transcriptomic divergence across mammals. (A) Total branch lengths of rodent DRG neuron orthotype gene expression trees (sum of transcriptomic distance/divergence values; see Methods). Box plots show the median (central value), upper and lower quartile (box limits), and 95% confidence intervals (whiskers) for 1,000 bootstrap replicates. Blue box highlights the three polymodal C-fiber (C-M/H) neuron orthotypes with the highest transcriptomic divergence values. (B) Rodent DRG neuron orthotype gene expression phylogenetic trees (see Methods). Neighbor-joining trees are generated using pairwise gene expression distances (1− ρ, Spearman’s correlation coefficient) between orthotypes both within and across species. Tree sizes shown here are normalized within orthotype. Bootstrap values (proportions of replicate trees with the same branching pattern as in the majority-rule consensus tree shown) lower than 99% are indicated by colored circles at the corresponding nodes. Orthotype gene expression phylogenetic trees largely do not recapitulate known phylogenetic relationships between species, but the invariability of intra-species replicate pairing demonstrates consistent gene expression patterns within orthotype across species.

**Fig. S12.**
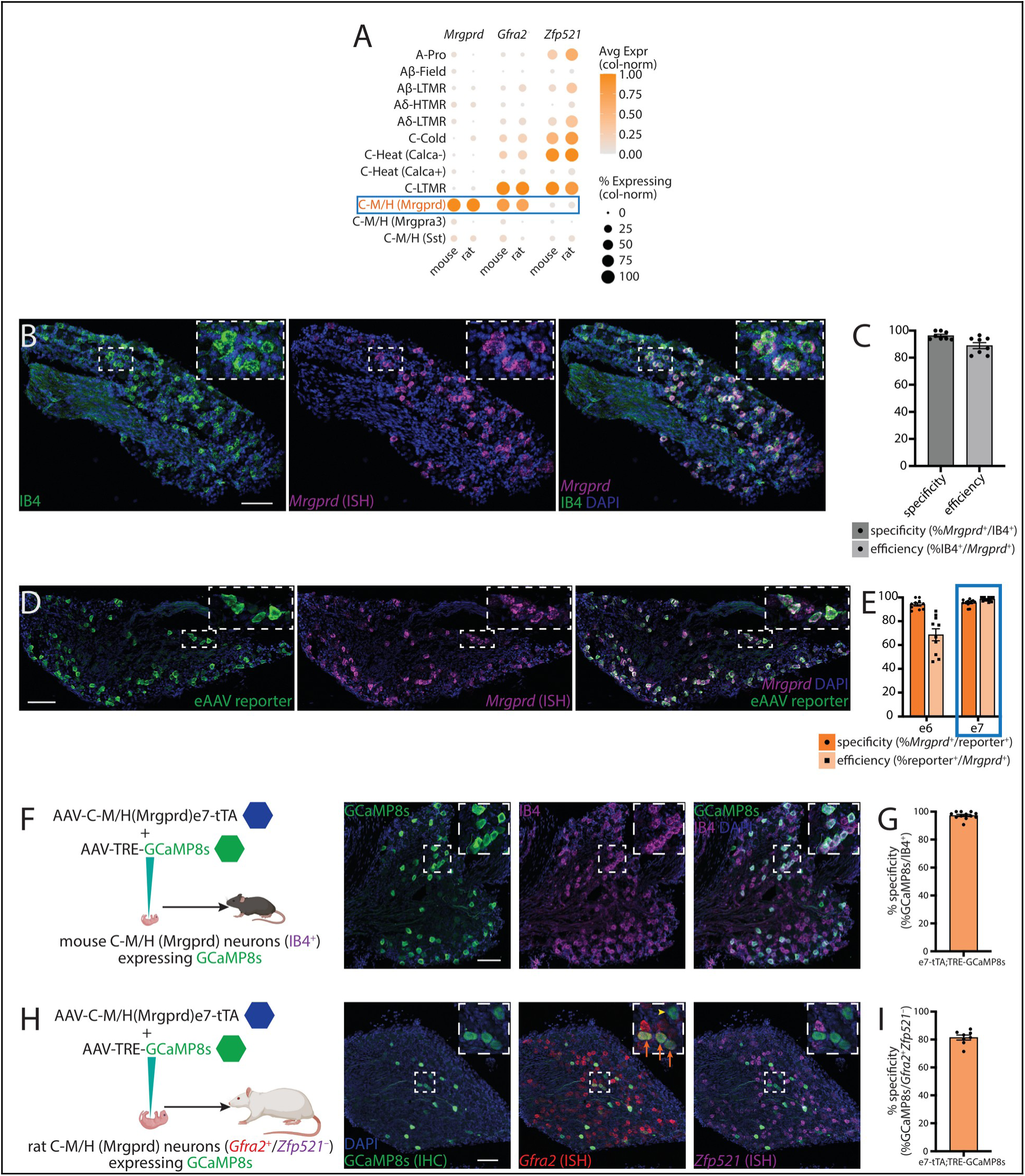
Additional validation of C-M/H (Mrgprd) neuron eAAV targeting in mice and rats. (A) Dot plot showing expression of *Mrgprd*, *Gfra2*, and *Zfp521* in mouse and rat DRG neuron orthotypes. C-M/H (Mrgprd) neurons are the only DRG neuron orthotype that expresses high *Gfra2* but no appreciable *Zfp521*. (**B** and **C**) Histology showing that IB4-binding is a property of C-M/H (Mrgprd) neurons in mice, marked by *Mrgprd in situ* hybridization signal. Fluorescence images (B). Quantification of specificity and efficiency (C); mean ± SEM. (**D** and **E**) Joint *in situ* hybridization-immunohistochemical validation of eAAV payload reporter expression in mouse C-M/H (Mrgprd) neurons (*Mrgprd*^+^). Fluorescence images (D). Quantification of specificity and efficiency (E); mean ± SEM. Blue box indicates eAAV selected for use in further experiments. (**F** to **I**) Schematics illustrating viral strategy for high GCaMP8s expression in mouse (F) and rat (H) C-M/H (Mrgprd) neurons. Fluorescence images displaying GCaMP8s expression in mouse (F, IB4^+^) and rat (H, *Gfra2*^+^/*Zfp521*^–^) C-M/H (Mrgprd) neurons. Quantification of specificity of GCaMP8s expression in mouse (G) and rat (I) C-M/H (Mrgprd) neurons; mean ± SEM. Scale bars, 100 µm.

**Fig. S13.**
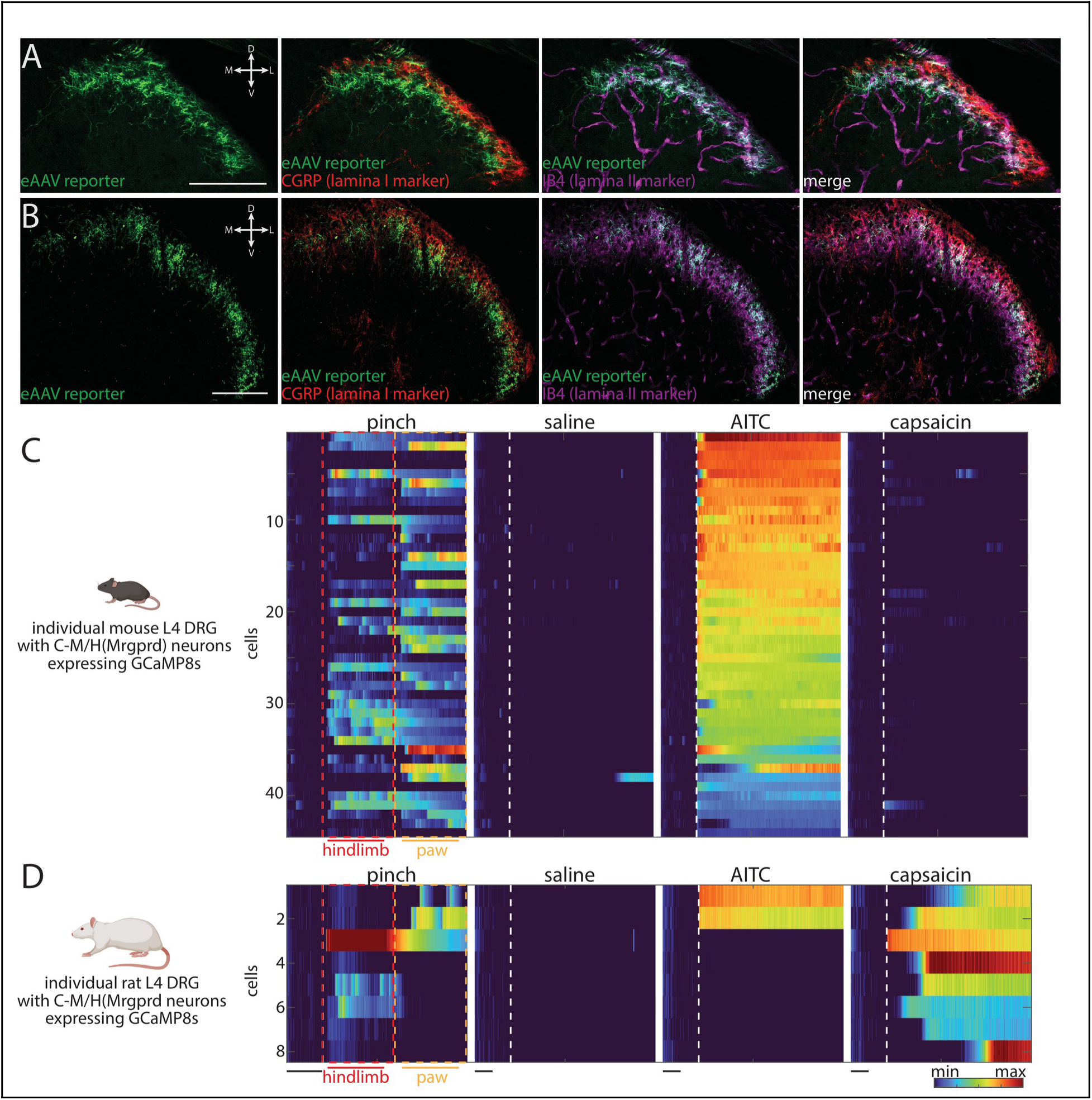
Additional characterization of eAAV-mediated C-M/H (Mrgprd) neuron anatomy and physiology in mice and rats. (**A** and **B**) eAAV-mediated C-M/H (Mrgprd) neuron labeling in mice (A) and rats (B) reveals central terminals targeting spinal cord dorsal horn lamina II marked by overlap with IB4-binding signal. Scale bars, 100 µm. See also fig. S6G. (**C** and **D**) Example Ca^2+^ imaging results from single mouse (C) and rat (D) DRGs displaying responses to pinch, vehicle control (saline), and chemical stimuli. Dotted white lines, stimulus onset. Colored bar, fluorescence (ΔF/F). Black bars, 10 seconds.

**Fig. S14.**
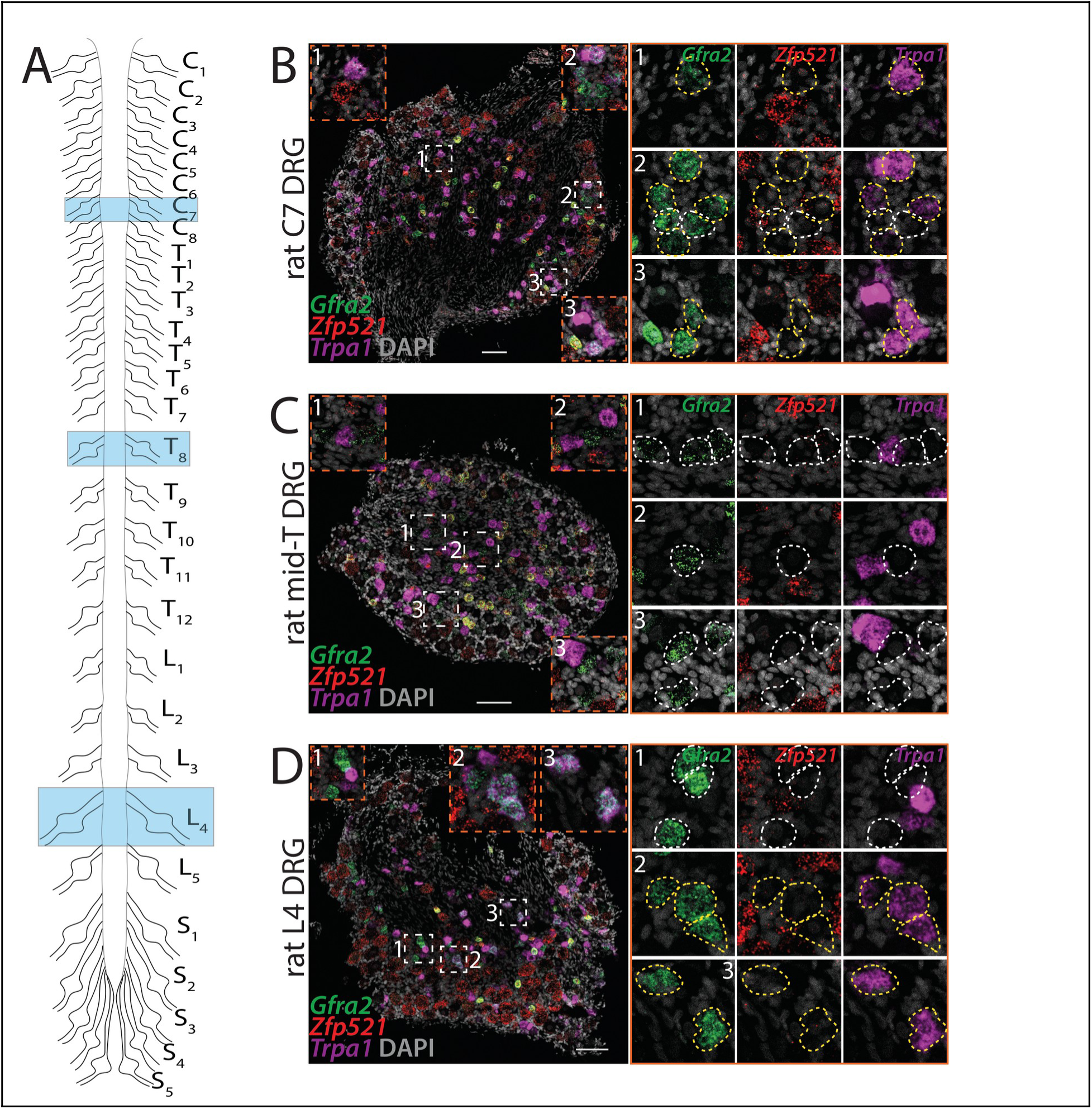
Histological identification of rat C-M/H (Mrgprd) neuron paratypes across axial levels. (A) Schematic illustrating DRGs taken of different axial levels. Note that cervical 7 (C7) and lumbar 4 (L4) axial level DRGs innervate distal limbs while mid-thoracic (mid-T, e.g., T8) axial level DRGs do not (see also Fig. 6C). (**B** to **D**) *In situ* hybridization images showing rat C-M/H (Mrgprd)_1_ (*Trpa1*^–^) and C-M/H (Mrgprd)_2_ (*Trpa1*^+^) neuron paratypes across C7, mid-T, and L4 DRGs. C-M/H (Mrgprd) neurons are *Gfra2*^+^/*Zfp521*^–^ (fig. S12A) and are indicated in the insets (right) by dotted white (*Trpa1*^–^) and yellow (*Trpa1*^+^) outlines. See quantification in Fig. 6, D to F. Scale bars, 100 µm.

**Fig. S15.**
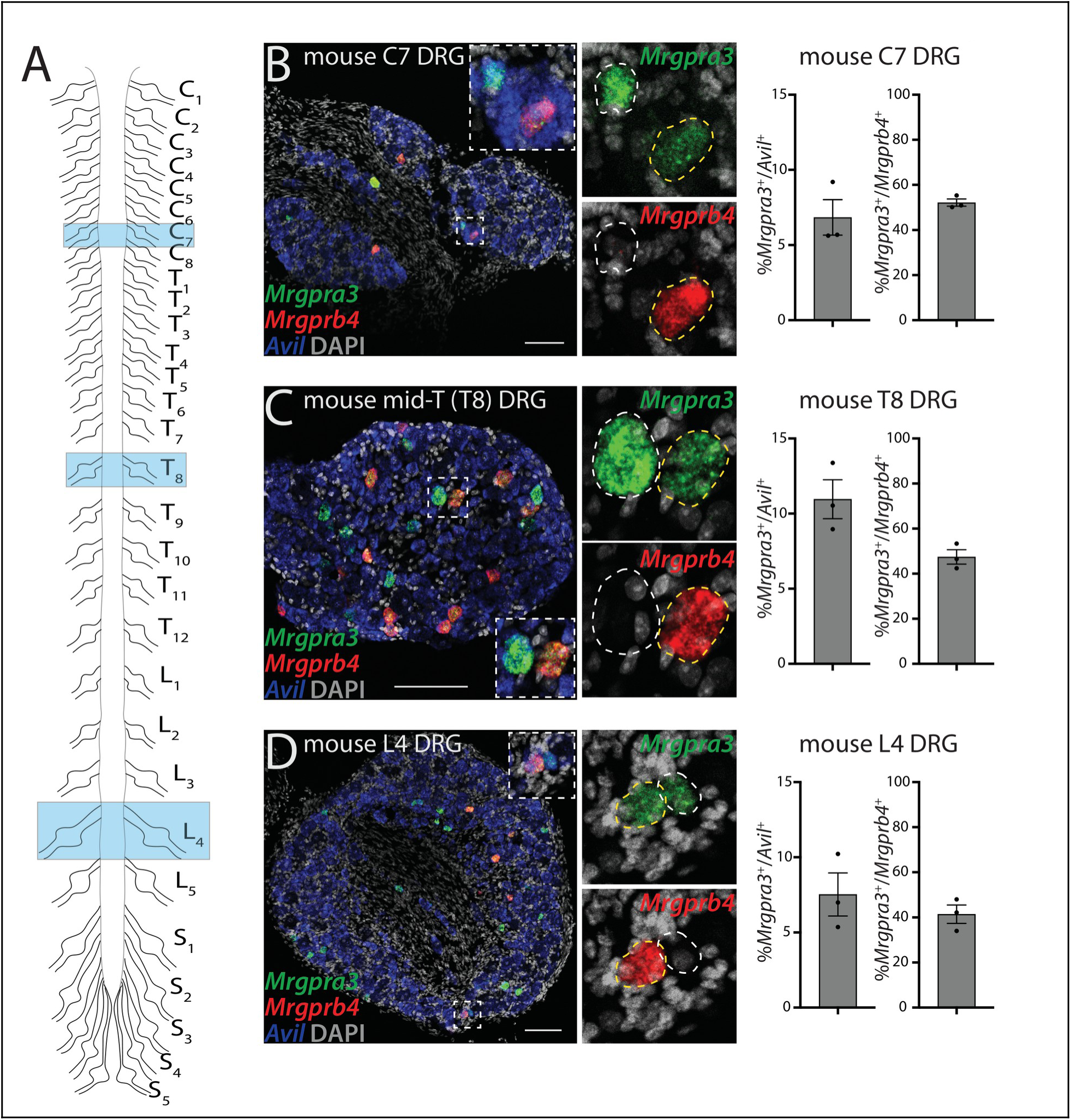
Histological identification of mouse C-M/H (Mrgpra3) neuron paratypes across axial levels. (A) Schematic illustrating DRGs taken of different axial levels. Note that cervical 7 (C7) and lumbar 4 (L4) axial level DRGs innervate distal limbs while mid-thoracic (mid-T, e.g., T8) axial level DRGs do not (see also Fig. 6C). (**B** to **D**) *In situ* hybridization images (left) and percentages (right) of mouse C-M/H (Mrgpra3)_1_ (only *Mrgpra3*^+^) and C-M/H (Mrgpra3)_2_ (*Mrgpra3*^+^*Mrgprb4*^+^) neuron paratypes across C7, mid-T (T8), and L4 DRGs; mean ± SEM. Scale bars, 100 µm.

**Fig. S16.**
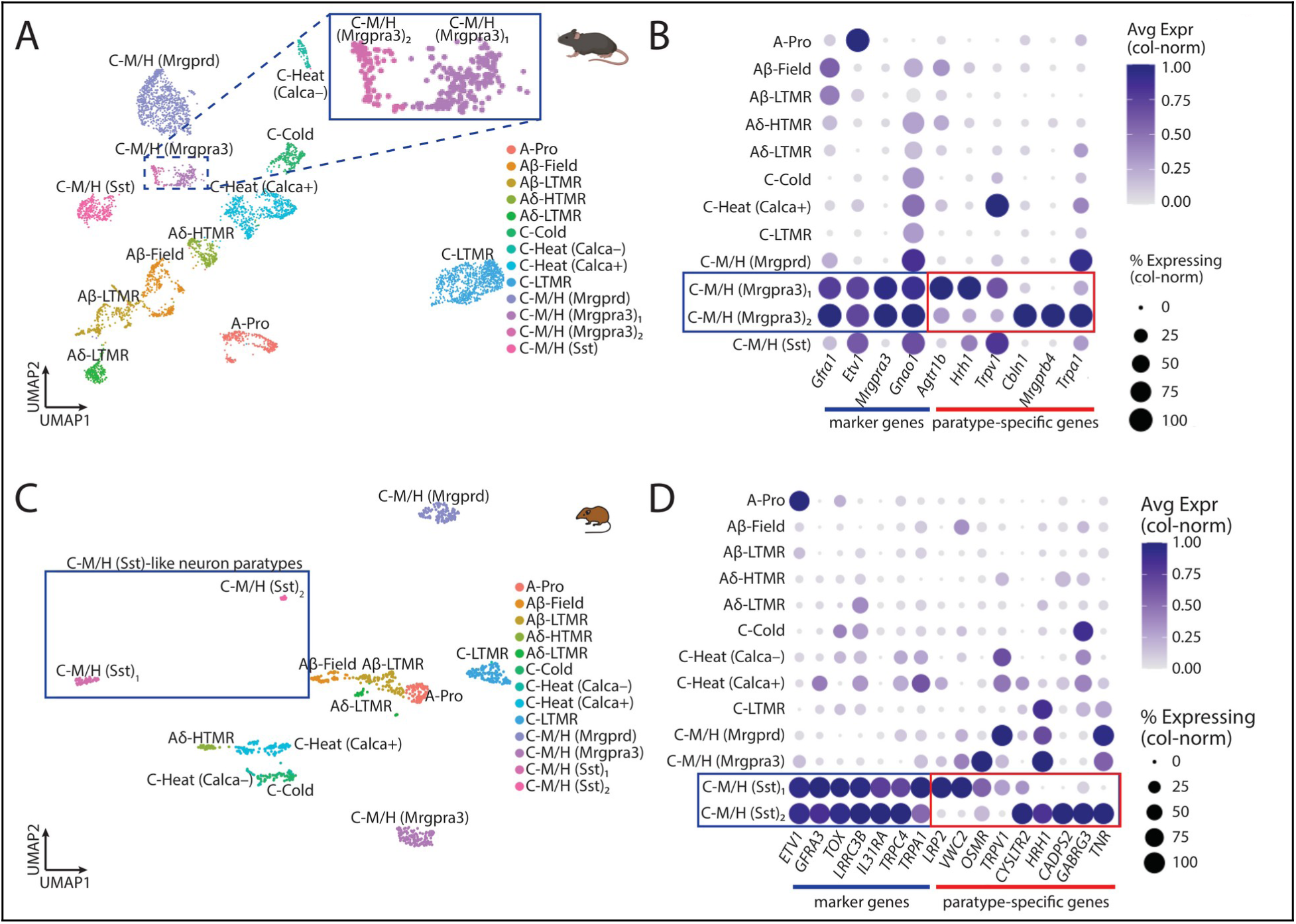
Bioinformatic identification and characterization of mouse C-M/H (Mrgpra3) and Etruscan shrew C-M/H (Sst) neuron paratypes. (A) UMAP embedding of snRNA-seq data from an individual mouse sample replicate. Colors indicate somatosensory neuron orthotype identities. Inset shows splitting of C-M/H (Mrgpra3) neuron orthotype into two putative paratypes. (B) Dot plot showing expression of common, orthotype-defining marker genes (blue line/box) and paratype-defining positive marker genes (red line/box) of mouse C-M/H (Mrgpra3) neuron paratypes. (C) UMAP embedding of snRNA-seq data from an individual shrew sample replicate. Colors indicate somatosensory neuron orthotype identities. Box shows splitting of C-M/H (Sst) neuron orthotype into two putative paratypes. (D) Dot plot showing expression of common, orthotype-defining marker genes (blue line/box) as well as paratype-defining positive marker genes (red line/box) of shrew C-M/H (Sst) neuron paratypes.

